# Ca^2+^ signaling in astrocytes is sleep-wake state specific and modulates sleep

**DOI:** 10.1101/750281

**Authors:** Laura Bojarskaite, Daniel M. Bjørnstad, Klas H. Pettersen, Céline Cunen, Gudmund Horn Hermansen, Knut Sindre Åbjørsbråten, Rolf Sprengel, Koen Vervaeke, Wannan Tang, Rune Enger, Erlend A. Nagelhus

## Abstract

Astrocytic Ca^2+^ signaling has been intensively studied in health and disease but remains uncharacterized in sleep. Here, we employed a novel activity-based algorithm to assess astrocytic Ca^2+^ signals in the barrel cortex of awake and naturally sleeping mice while monitoring neuronal Ca^2+^ activity, brain rhythms and behavior. We discovered that Ca^2+^ signaling in astrocytes exhibits distinct features across the sleep-wake cycle and is reduced in sleep compared to wakefulness. Moreover, an increase in astrocytic Ca^2+^ signaling precedes transitions from slow-wave sleep to wakefulness, with a peak upon awakening exceeding the levels during whisking and locomotion. Genetic ablation of a key astrocytic Ca^2+^ signaling pathway resulted in fragmentation of slow-wave sleep, yet increased the frequency of sleep spindles. Our findings suggest a role for astrocytic Ca^2+^ signaling in modulating sleep.

## INTRODUCTION

We spend approximately one third of our lives sleeping. The purpose of sleep is one of the greatest unsolved mysteries in biology, although it is increasingly clear that sleep is important for higher-order cognitive functions and restoration (Rasch and Born, 2013). Still, the nightlife of the housekeepers of the brain – the astrocytes – is poorly characterized. There is evidence that astrocytes regulate sleep drive (Halassa et al., 2009), promote sleep-dependent brain waste clearance (Iliff et al., 2012; Xie et al., 2013) and facilitate cortical oscillations that are important for learning and memory (Poskanzer and Yuste, 2016; Szabó et al., 2017), but the signaling mechanisms that astrocytes employ to mediate these sleep-dependent functions remain elusive.

Astrocytic Ca^2+^ signals are considered to orchestrate neuronal circuits by regulating the extracellular ion concentration and promoting the release of signaling substances (Bazargani and Attwell, 2016; Dallérac et al., 2018; Verkhratsky and Nedergaard, 2018; Volterra et al., 2014). In addition, astrocytes not only sense local synaptic activity (Haustein et al., 2014; Panatier et al., 2011), but also respond with Ca^2+^ signals to neuromodulators (Araque et al., 2002; Paukert et al., 2014; Takata et al., 2011) that are involved in sleep-wake state regulation (Lee and Dan, 2012; Scammell et al., 2017). Whereas attempts have been made to image astrocytic Ca^2+^ signals under anesthetic conditions mimicking sleep (Poskanzer and Yuste, 2016; Szabó et al., 2017), astrocytic Ca^2+^ signaling in natural sleep has to the best of our knowledge not yet been characterized.

Recent advances in optical imaging and genetically encoded activity sensors have enabled high resolution imaging of astrocytes in unanesthetized mice, revealing an exceedingly rich repertoire of astrocytic Ca^2+^ signals (Bindocci et al., 2017; Srinivasan et al., 2015; Stobart et al., 2018). Conventional tools for analysis of astrocytic Ca^2+^ signals are based on static regions-of-interest (ROIs) placed over morphologically distinct compartments. However, studies in the last few years indicate a highly complex and dynamic spatiotemporal nature of astrocytic Ca^2+^ signals, which is not captured by the ROI analyses (Bindocci et al., 2017; Srinivasan et al., 2015). New standardized methods for astrocyte Ca^2+^ analyses are warranted.

We set out to explore astrocytic Ca^2+^ signaling in mice during natural head-fixed sleep. We employed dual-color two-photon Ca^2+^ imaging to observe cortical astrocytes and neurons simultaneously, combined with electrocorticography (ECoG), electromyography (EMG) and behavioral monitoring. Using a novel, automated activity-based analysis tool we reveal a new level of complexity of astrocytic Ca^2+^ signals. We show that Ca^2+^ signaling in astrocytes is reduced during sleep compared to wakefulness and exhibits distinct characteristics across different sleep states. We demonstrate that an increase in astrocytic Ca^2+^ signaling precedes transition from slow-wave sleep (SWS) – but not rapid eye movement (REM) sleep – to wakefulness. Finally, we demonstrate that the inositol triphosphate (IP_3_)-mediated astrocytic Ca^2+^ signaling pathway modulates sleep states, brain rhythms and sleep spindle dynamics. Our data suggest an essential role for astrocytic Ca^2+^ signaling in modulating sleep.

## RESULTS

### Two-photon imaging of Ca^2+^ signals in cortical astrocytes and neurons in awake and naturally sleeping mice

To explore the characteristics of Ca^2+^ signaling in astrocytes and neurons across sleep-wake states, we employed dual-color two-photon Ca^2+^ imaging of neurons and astrocytes in layer II/III of the mouse barrel cortex by combining the green Ca^2+^ indicator GCaMP6f in astrocytes with the red Ca^2+^ indicator jRGECO1a in neurons (Figure 1A). The *glial fibrillary acidic protein* (*GFAP*) and the human *synapsin1 (SYN)* promoters were used to target astrocytes and neurons, respectively. In accordance with recent reports underscoring the importance of high image acquisition rates for capturing fast populations of astrocytic Ca^2+^ events (Bindocci et al., 2017), all our imaging was performed at a frame rate of 30 Hz. Concomitantly, we recorded mouse behavior with an infrared (IR) sensitive camera, electrocorticography (ECoG) and EMG for classification of sleep-wake states (Figure 1A). The transduced *SYN*-jRGECO1a and *GFAP*-GCaMP6f specifically labeled neurons and astrocytes without inducing astrogliosis or microglia activation (Figure S1). We identified three different behavioral states of wakefulness by analyzing mouse movements on the IR video footage (Figures 1B, 1E and 1F): voluntary locomotion, spontaneous whisking and quiet wakefulness. Since locomotion in mice and rats is tightly associated with natural whisking (Sofroniew et al., 2014), our locomotion behavioral state comprises both movement and whisking (Figure 1B). Using standard criteria on ECoG and EMG (Niethard et al., 2016; Oishi et al., 2016; Seibt et al., 2017) signals we identified three sleep states (Figures 1C–F): non-rapid eye movement (NREM) sleep, intermediate state (IS) sleep and REM sleep. NREM sleep and IS sleep are sub-states of SWS. IS sleep is a transitional state from NREM to REM sleep, found at the end of a NREM episode and characterized by an increase in sleep spindle frequency, increase in sigma (10– 16 Hz) and theta (5–9 Hz) ECoG power and a concomitant decrease in delta (0.5–4 Hz) oscillations (Seibt et al., 2017) (Figure 1C–E). We excluded microarousals in SWS (Watson et al., 2015) since we aimed to characterize true sleep astrocytic Ca^2+^ signals. Head-fixed sleeping mice exhibited normal spectral ECoG properties and sleep architecture (Figures 1E and 6A–C) (Cao et al., 2013; Foley et al., 2017; Oishi et al., 2016; Seibt et al., 2017). To capture the high spatiotemporal complexity of astrocytic Ca^2+^ signaling, we quantified astrocytic Ca^2+^ activity using a novel automated activity-based analysis tool (Figure S2). The algorithm comprises three-dimensional filtering and noise-based thresholding on individual pixels over time to detect fluorescent events. Connecting adjacent active pixels in space and time results in ‘regions-of-activity’ (ROAs) that can subsequently be combined with conventional, manually drawn ROIs or analyzed separately. The specificity of the algorithm was assured by testing the ROA method on time series from control mice expressing a Ca^2+^ insensitive fluorescent indicator (enhanced green fluorescent protein, eGFP) in cortical astrocytes (Figure S3). We then compared the characteristics of Ca^2+^ event detection with ROI and ROA analyses (Figure S4). Ca^2+^ signaling in astrocytic somata and processes appeared to be much more complex than captured by static ROIs. Ca^2+^ signals were highly heterogeneous, ranging from small events in astrocytic subdomains to events involving all astrocyte domains in a field-of-view (FOV; Figures S4A and S4B). Notably, small, low amplitude events in microdomains remained undetected with ROI analysis due to low ratio of active area to the total area outlined by a manual ROI (Figure S4C and S4D), resulting in up to 60% fewer detected events (Figure S4E and S4F). These data indicate that conventional static morphology-based ROIs fail to reveal the complexity of astrocyte Ca^2+^ signaling. In all, 13 hours of wakefulness and over 15 hours of natural sleep (7 h NREM, 5 h IS, 3 h REM) (Figure 1E) in 6 wild type (WT) mice were analyzed. Representative wakefulness and sleep trials are shown in Figures S5 and S6.

**Figure 1.**
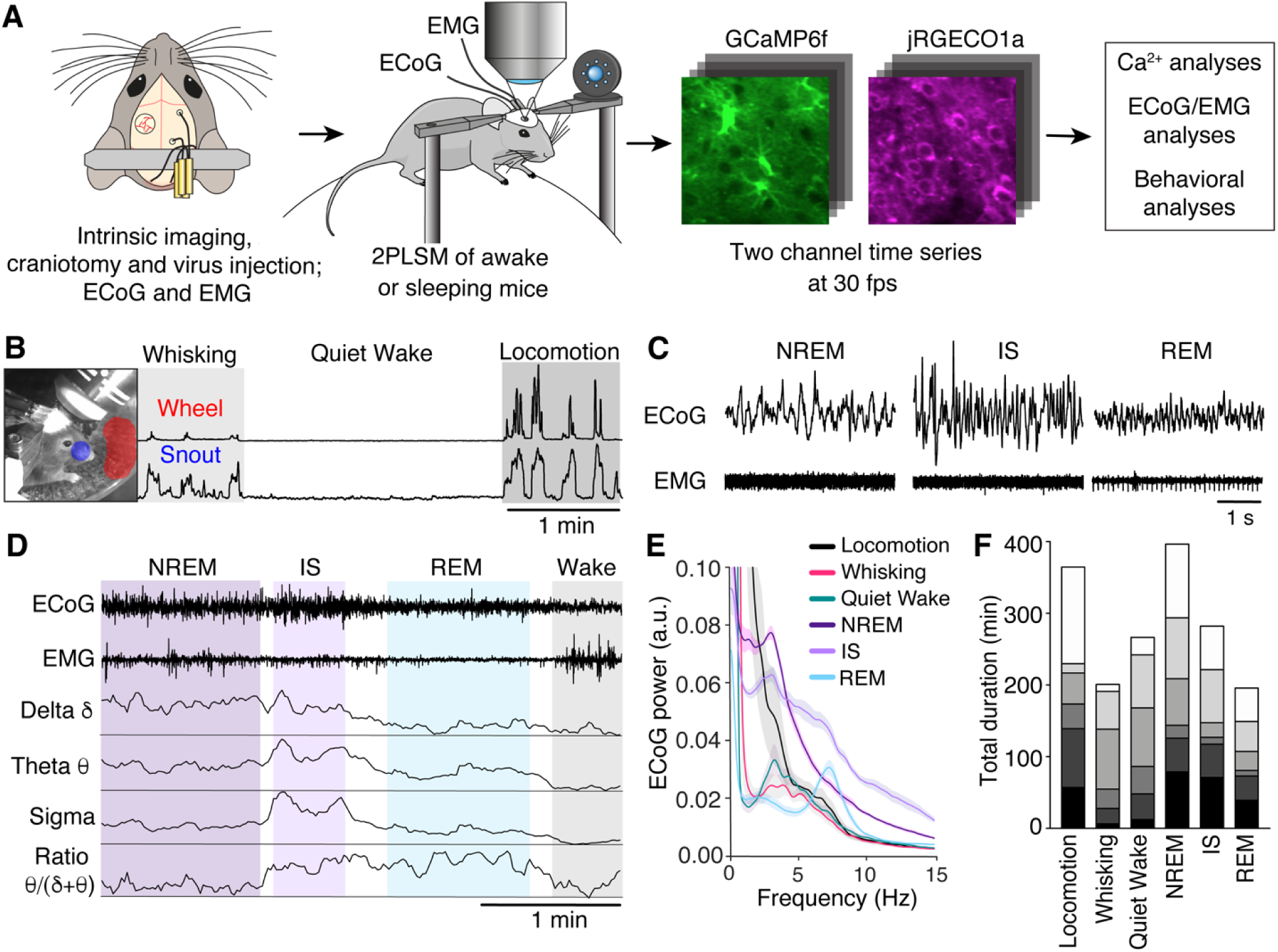
Two-photon dual-color imaging of Ca^2+^ signals of astrocytes and neurons in awake and naturally sleeping mice. (A) Experimental setup. A craniotomy exposing the barrel cortex was made, and a mixture of rAAV-*GFAP*-GCaMP6f and rAAV-*SYN*-jRGECO1a were injected for visualization of astrocytic and neuronal Ca^2+^ signals, respectively. ECoG and EMG electrodes were implanted, and in combination with IR surveillance video allowed classification of the sleep-wake state. (B) Wakefulness was separated into locomotion, whisking and quiet wakefulness based on movement of the mouse’s snout and of the running wheel detected in the surveillance video. (C) Representative ECoG and EMG recordings of NREM, IS and REM sleep states. (D) Sleep is categorized into NREM, IS and REM sleep states based on the delta (δ), theta (θ) and sigma (σ) frequency band power, EMG and theta to delta frequency band ratio. (E) Normalized ECoG power density spectrum of sleep-wake states, n = 6 mice, average ± SEM. (F) Total duration of each wake-sleep state from every mouse. Individual mice are indicated as separate blocks in bars, n = 6 mice. See also Figures S2–S7.

### Astrocytic Ca^2+^ signaling is reduced during sleep and is sleep state specific

We used the ROA analysis to explore astrocytic Ca^2+^ signaling during wakefulness and natural sleep (Figure 2). We discovered a broad repertoire of astrocytic Ca^2+^ signals across the sleep-wake cycle. The spatial extent of ROAs ranged from ∼0.9 μm^2^ (lower detection limit) to the full FOV (Figure S7A), whereas the duration of the events ranged from 0.1 s to 60 s (Figure S7B). The majority of astrocytic Ca^2+^ events were small and short-lasting (∼80% events < 10 μm^2^ and < 1s). ROA event size and duration were positively correlated, with the larger events lasting longer (Supplementary Figure S7D). The largest events were detected almost exclusively during locomotion (Figures 2A and S5). Such synchronized Ca^2+^ events in astrocytes started at multiple foci and eventually merged to cover almost the entire FOV, similar to what has previously been reported (Bindocci et al., 2017).

**Figure 2.**
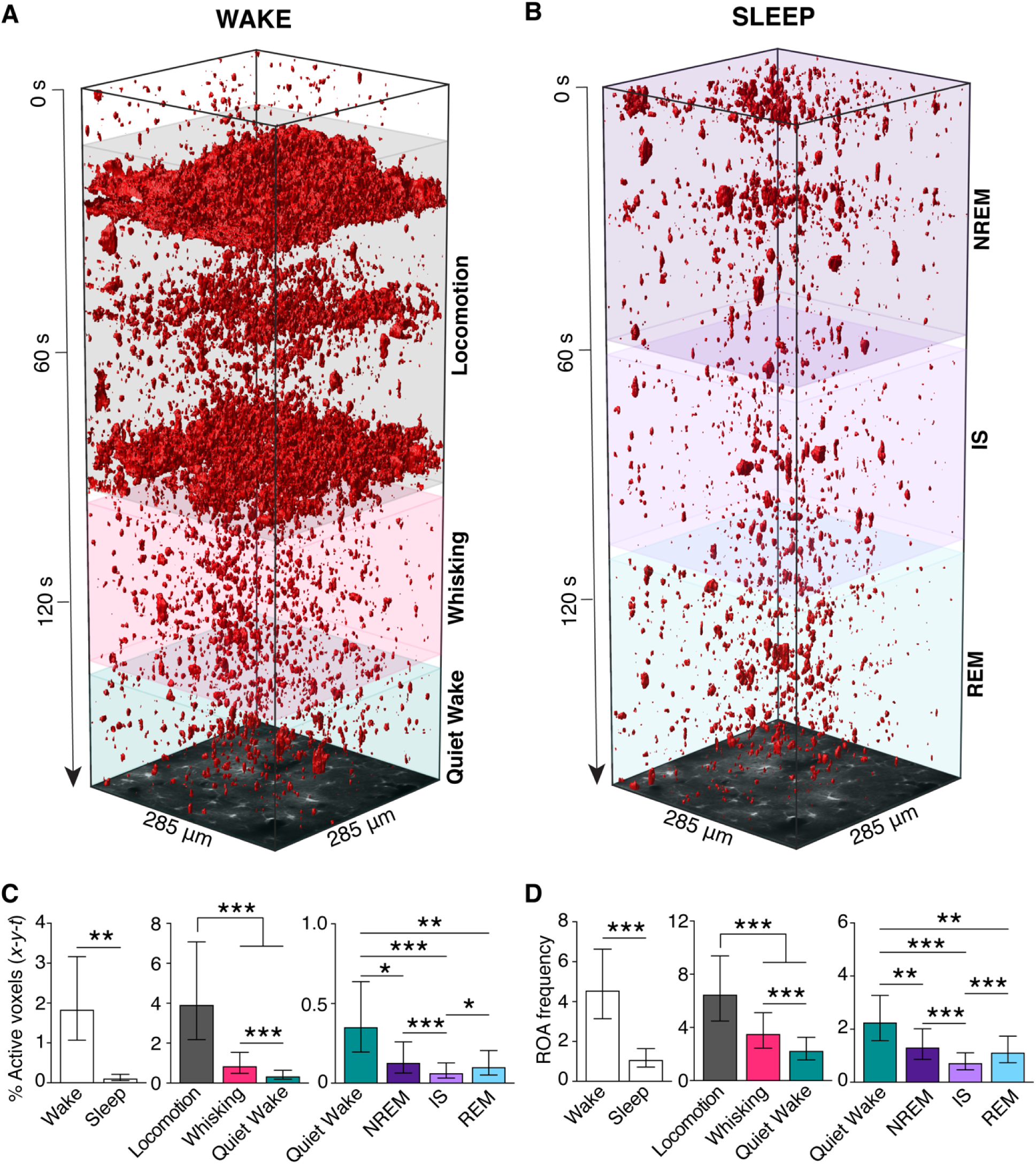
Astrocytic Ca^2+^ signaling is brain state specific and reduced during sleep. (A and B) Representative *x-y-t* rendering of ROAs during wakefulness (A) and sleep (B). (C) Percentage of active voxels (*x-y-t*) during overall wakefulness and overall sleep (*left*), during locomotion, whisking and quiet wakefulness (*middle*), and during NREM, IS and REM sleep as compared to quiet wakefulness (*right*). (D) Same as (C) but for ROA frequency expressed as number of ROAs per 100 μm^2^ per minute. Data represented as estimates ± SEM, n = 6 mice, 283 trials, *p<0.05, **p<0.005, ***p<0.0005. For details on statistical analyses, see Methods. See also Figures S5–S8.

We then quantified astrocytic Ca^2+^ activity across sleep-wake states. We first compared Ca^2+^ activity between overall wakefulness and overall sleep. Astrocytic Ca^2+^ activity was reduced during sleep compared to wakefulness with a 94% reduction in the mean percent of active voxels (*x-y-t*) and a 77% reduction in the frequency of ROAs (Figures 2C and 2D, left). Since wakefulness encompasses a spectrum of sub-states that serve distinct, perceptual and behavioral functions (McGinley et al., 2015), we investigated astrocyte Ca^2+^ signaling across the different states of wakefulness. Voluntary locomotion and spontaneous whisking increased the percentage of active voxels (11-fold during locomotion, 2.5-fold during whisking) and increased ROA frequency (3-fold during locomotion, 1.5-fold during whisking) compared to quiet wakefulness (Figures 2C and 2D, middle). We chose the resting state – quiet wakefulness – for comparison with specific sleep states. Astrocytic Ca^2+^ activity was reduced in all sleep states compared to quiet wakefulness as measured by both the percentage of active voxels (*x-y-t*) and ROA frequency (Figures 2C and 2D, right). Moreover, astrocytic Ca^2+^ activity was sleep-state specific: The percentage of active voxels (*x-y-t*) and ROA frequency was lower during IS sleep than during NREM or REM sleep (Figures 2C and 2D, right). Conventional ROI analyses of Ca^2+^ signals in astrocytes showed similar tendencies across sleep-wake states, but failed to capture both the difference between quiet wakefulness and sleep and the differences between the sleep states (Figures S8). Taking advantage of this novel activity-based ROA analysis, we provide the first demonstration that astrocytic Ca^2+^ signaling is reduced during sleep compared to wakefulness, and is sleep-state specific.

### Astrocytic Ca^2+^ signals are most frequent in processes during all sleep-wake states

Since Ca^2+^ transients in astrocytic somata and processes may have different underpinning and function (Khakh and Sofroniew, 2015; Srinivasan et al., 2015) we investigated the spatial distribution of the Ca^2+^ signals across sleep-wake states (Figure 3A). Activity maps indicated that the astrocytic Ca^2+^ activity was concentrated in their processes in neuropil for all brain states. To quantify the spatial distribution of the Ca^2+^ signals we ran the ROA algorithm within manually drawn ROIs over astrocytic somata and neuropil (Figure 3B). Despite that astrocytic processes occupy only 8.5% of barrel cortex neuropil volume in adult mice (Genoud et al., 2006), the frequency of astrocytic Ca^2+^ signals was higher in neuropil ROIs than in astrocytic somata ROIs across all sleep-wake states (Figure 3C). Still, both astrocytic somata and processes displayed similar magnitude and direction of changes in Ca^2+^ signaling across sleep-wake states (Figure 3C). To conclude, astrocytic Ca^2+^ signals are more frequent in processes than somata in all sleep-wake states.

**Figure 3.**
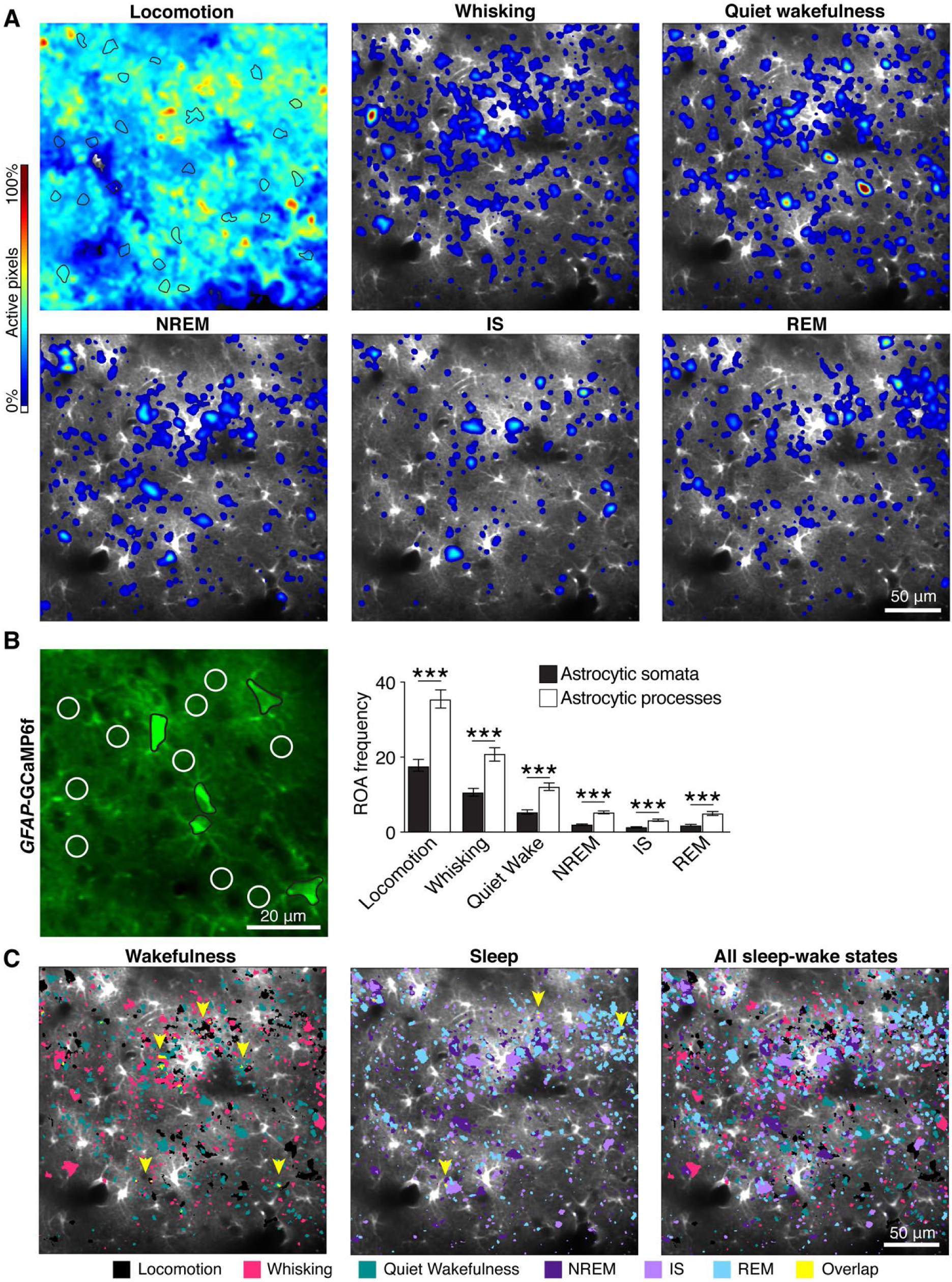
Astrocytic Ca^2+^ signals are most frequent in processes during all sleep-wake states. (A) Representative ROA frequency heatmaps showing the localization of Ca^2+^ signals during single episodes of locomotion, whisking, quiet wakefulness, NREM sleep, IS sleep and REM sleep states. Astrocytic somata outlined in black in the locomotion image. (B) *Left*: Representative image of *GFAP*-GCaMP6f fluorescence in astrocytes with ROIs over astrocytic somata (black) and neuropil, containing astrocytic processes (white circles of 10 μm in diameter). *Right*: ROA frequency, expressed as number of ROAs per 100 μm^2^ per minute, in astrocyte somata and processes from ROIs shown in left across sleep-wake states. Data represented as estimates ± SEM, n = 6 mice, 247 trials, *p<0.05, **p<0.005, ***p<0.0005. For details on statistical analyses, see STAR Methods. (C) Representative Ca^2+^ activity maps displaying the top 5% most active pixels in locomotion (black), whisking (pink), quiet wakefulness (green), NREM sleep (dark purple), IS sleep (light purple), and REM sleep (light blue). Yellow areas indicated by yellow arrows are areas of overlapping Ca^2+^ hot spots across states indicated above the images.

To study the extent to which astrocytes maintained areas with particularly high Ca^2+^ activity (‘hotspots’), we created maps displaying only the top 5% most active pixels within a given sleep-wake state and assessed the overlap between Ca^2+^ hotspots across sleep-wake states (Figure 3D). Astrocytic Ca^2+^ hotspots typically shifted location between states, in both wakefulness and sleep. Only a small fraction of the hotspot area persisted across all wakefulness states (3.3 ± 0.3 %, n = 66 FOVs), across all sleep states (2.1 ± 0.3 %, n = 82 FOVs), or across all sleep-wake states (0.2 ± 0.05 %, n = 60 FOVs). Thus, our analysis indicated poor spatial stability of Ca^2+^ hotspots across sleep-wake states.

### Astrocytic Ca^2+^ signals increase prominently upon transitions from sleep to wakefulness and precede awakenings from SWS, but not from REM sleep

We observed that astrocytic Ca^2+^ activity was not homogeneously distributed within a given brain state, but rather was concentrated at transitions from one state to another (Figures S5 and S6, red squares). To explore the relationship between state transitions and astrocytic Ca^2+^ activity, we plotted ROA frequency aligned to the beginning of states, i.e. from quiet wakefulness to locomotion or whisking, and from either NREM, IS or REM sleep to wakefulness (Figure 4A). The onset of locomotion and whisking was automatically defined as a movement of the whisker or wheel in the surveillance video, whereas wakefulness onset during sleep-to-wake transitions was manually determined as the first sign of ECoG desynchronization (Takahashi et al., 2010) (Figure 4B). Astrocytic Ca^2+^ events were strongly clustered around specific brain state transitions (Figure 4A and 4C–D). During transitions from quiet wakefulness to locomotion or whisking, astrocytic Ca^2+^ signaling was delayed compared to the transition in the ECoG signal, muscle activity, and mouse movement in most cases (Figures 4A and 4C, 4E). The average onset delay of astrocytic Ca^2+^ did not differ for transitions to locomotion or whisking, but the peak Ca^2+^ response in relation to the transition was larger for locomotion (Figures 4F–G). We then investigated the relationship between astrocytic Ca^2+^ activity and the transition from sleep to wakefulness (Figures 4A–B and 4D). To our surprise, astrocytic Ca^2+^ signals preceded the awakenings from SWS (NREM and IS). We observed a prominent increase in astrocytic Ca^2+^ signal frequency 1 – 2 seconds before the shift in ECoG, EMG and mouse movement in 60% of NREM sleep to wakefulness and 72% of IS sleep to wakefulness transitions (Figures 4A, 4D–F). As astrocytic Ca^2+^ signaling did not precede the transition from NREM or IS sleep to wakefulness in all cases, we investigated if ECoG power or sleep bout duration could determine the temporal profile of astrocytic Ca^2+^ signal onset (Figure S9). We found that high delta ECoG power was associated with earlier astrocytic Ca^2+^ onset in NREM sleep, but not IS sleep (Figures S9A). Moreover, shorter IS bout duration was associated with earlier astrocytic Ca^2+^ onset in IS sleep, but no such trends were observed in NREM sleep (Figure S9D). We did not find correlation between astrocytic Ca^2+^ onset and ECoG power in the theta and sigma range in neither NREM nor IS sleep to wakefulness transition (Figures S9B and S9C).

**Figure 4.**
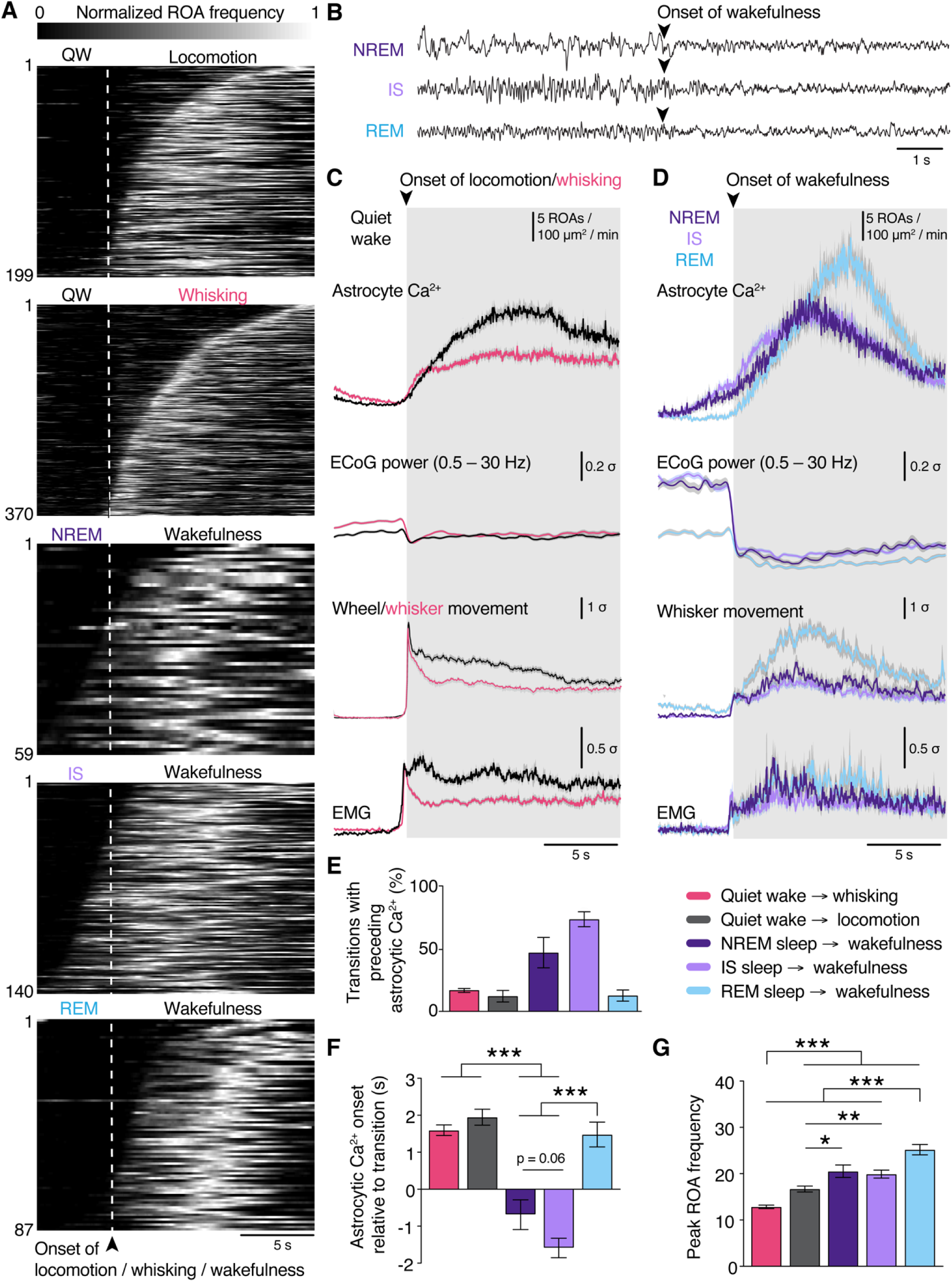
Astrocytic Ca^2+^ signals increase prominently upon awakening and precede transitions to wake from SWS, but not from REM sleep. (A) Temporal raster plots of normalized ROA frequency during the transitions (top to bottom): quiet wakefulness to locomotion (n = 199 transitions); quiet wakefulness to whisking (n = 370 transitions); NREM sleep to wakefulness (n = 59 transitions); IS sleep to wakefulness (n = 140 transitions); REM sleep to wakefulness (n = 87 transitions). (B) Examples of transitions from NREM, IS and REM sleep to wakefulness in the ECoG traces. (C) Mean time-course of (top to bottom) astrocytic Ca^2+^ signals and z-scores of ECoG power, wheel or whisker motion and EMG activation during transitions from quiet wakefulness to locomotion or whisking. Data represented as mean ± SEM, n = 6 mice (D) Same as (C) but for transitions from NREM, IS or REM sleep to wakefulness. (E) Percentage of transitions with preceding astrocytic Ca^2+^ signals. Data represented as mean ± SEM, n = 6 mice, number of samples for each transition described in (A). (F) Astrocytic Ca^2+^ signal onset relative to state transition. Values represent estimates ± SEM, n = 6 mice, number of samples for each transition described in (A). (G) Peak frequency of ROAs during state transitions. Values represent estimates ± SEM, n = 6 mice, number of samples for each transition described in (A). *p<0.05, **p<0.005, ***p<0.0005. See Methods for details on statistical analyses. See also Figures S9 and S10.

Waking up from REM sleep was more similar to state transitions of wakefulness as astrocytic responses were delayed compared to the transition (Figure 4A, 4D and 4F). The peak astrocytic Ca^2+^ response in the transition to wakefulness was larger when waking up from REM sleep compared to waking up from SWS or transitions from quiet wakefulness to whisking and locomotion (Figure 4G). On the contrary, transitioning from wakefulness to sleep was associated with a decrease in Ca^2+^ signals (Figure S10). To conclude, astrocytic Ca^2+^ signaling is enhanced upon awakenings and precedes transitions from SWS to wakefulness, suggesting that astrocytic Ca^2+^ signaling may play a role in modulating sleep-to-wake transitions.

### Astrocytic Ca^2+^ signaling in sleep differs from the overall local neuronal activity

Since astrocytic Ca^2+^ signaling could depend on local neuronal activity, we also assessed Ca^2+^ signals in neighboring neurons and the relationship between the signals in the two cell types. To capture neuronal Ca^2+^ activity, we measured jRGECO1a fluorescence in hand-drawn ROIs over neuronal somata and neuropil (Figure 5A). The frequency of Ca^2+^ signals in neuronal somata was lower during sleep compared to wakefulness, but the reduction was much less pronounced than that of astrocytes (Figures 5B and S11). We then examined the temporal relationship between astrocyte and neuron Ca^2+^ signals across sleep-wake states by calculating the onset of astrocytic Ca^2+^ signals in neuropil ROIs relative to Ca^2+^ events in neuronal soma ROIs (Figure 5C). During locomotion and spontaneous whisking, a population of astrocytic Ca^2+^ signals were tightly distributed around neuronal events (Figures 5C), indicating elevated astrocyte-neuron synchrony during sensory activation, similar to previous reports (Stobart et al., 2018). By contrast, during quiet wakefulness and all sleep states, astrocytic Ca^2+^ signals displayed a broad distribution of onset time differences relative to neighboring neurons (Figures 5C), suggesting that astrocytic Ca^2+^ is uncoupled from local neuronal activity during brain states that lack dense neuronal discharge.

**Figure 5.**
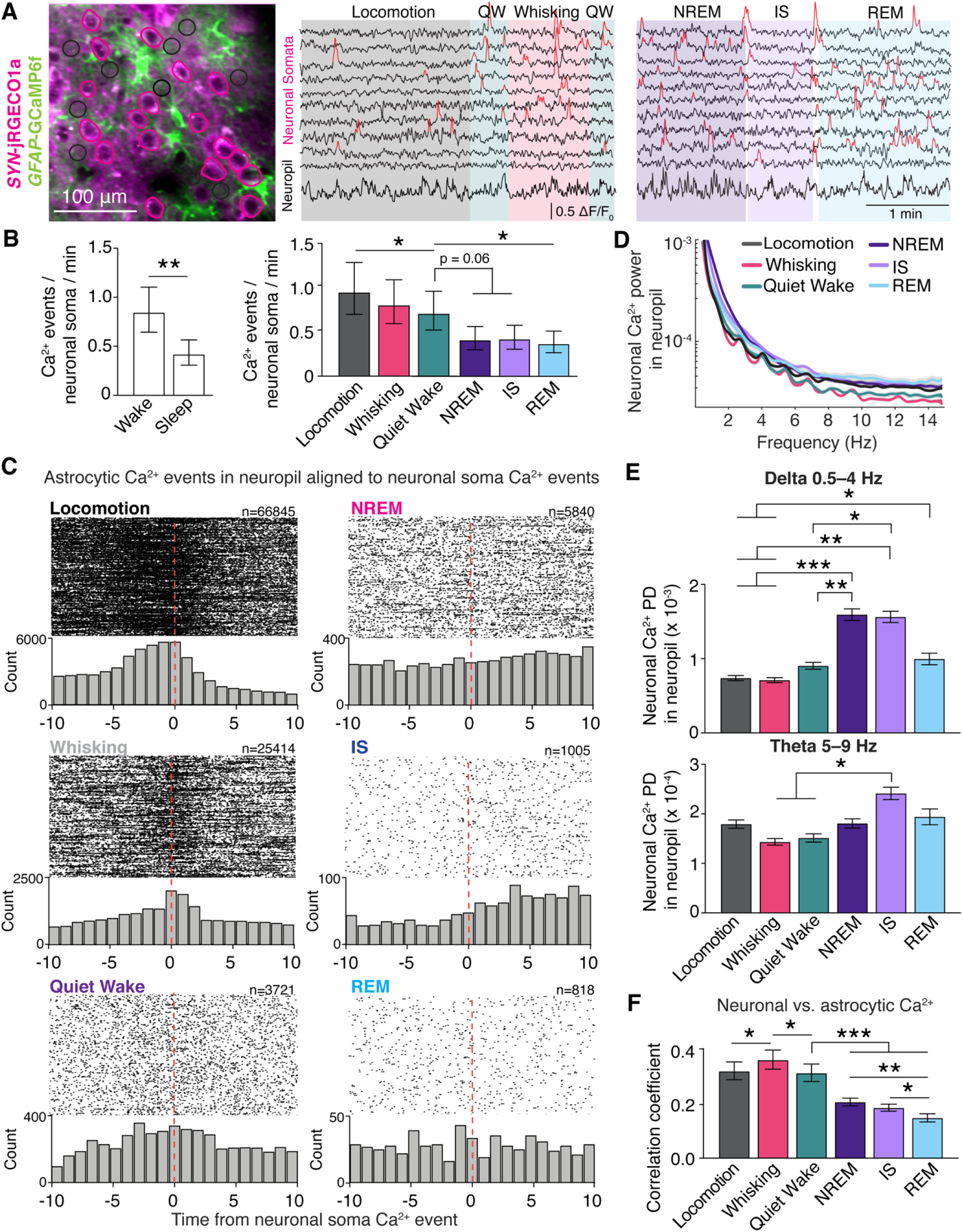
Astrocytic Ca^2+^ signaling in sleep differs from overall local neuronal activity. (A) *Left*: Representative image of *SYN*-jRGECO1a fluorescence in neurons and *GFAP*-GCaMP6f fluorescence in astrocytes, and ROIs over neuron somata (pink) and neuropil (black circles of 10 μm in diameter). *Right*: example ΔF/F_0_ Ca^2+^ traces from ROIs over neuron somata, smoothed by a 10-frame moving median filter, and neuropil in *SYN*-jRGECO1a channel across sleep-wake states. Detected Ca^2+^ peaks in neuronal somata are indicated in red. Neuropil Ca^2+^ signal calculated as average of all neuropil ROIs per FOV. (B) Frequency of Ca^2+^ signals in neuron somata during overall wakefulness and overall sleep (*left*), and across sleep-wake states (*right*). Data represented as estimates ± SEM, n = 6 mice, 92 trials. (C) Raster plots and histograms (1 s bins) of the onset time difference for astrocytic Ca^2+^ events in neuropil ROIs (black circular ROIs in A) relative to a Ca^2+^ event in neuron somata ROIs (pink ROIs over neuron somata in A). (D) Power spectra of the neuropil Ca^2+^ signal across sleep-wake states. Data represented as mean ± SEM, n = 6 mice, 92 trials. (E) Power density (PD) of the neuropil Ca^2+^ signal in delta 0.5–4 Hz and theta 5–9 Hz frequency bands across sleep-wake states. Data represented as estimates ± SEM, n = 6 mice, 92 trials. (F) Correlation coefficient between astrocytic Ca^2+^ events in neuropil, measured as number of ROAs per 100 μm^2^ per minute, and neuropil Ca^2+^ signal across sleep-wake states. Data represented as estimates ± SEM, n = 6 mice, 81 trials. *p<0.05, **p<0.005, ***p<0.0005. For details on statistical analyses, see Methods. See also Figure S11.

Next we analyzed neuronal Ca^2+^ signaling in neuropil, which is more representative of the input to the local neuronal circuitry (Kerr et al., 2005). By calculating the power spectrum of neuropil jRGECO1a fluorescence across sleep-wake states (Seibt et al., 2017), we detected increased power density (PD) in the low frequency ranges (delta 0.5–4 Hz) during SWS (NREM and IS sleep states) compared to all states of wakefulness, potentially representing slow network oscillations related to SWS (Niethard et al., 2018) (Figure 5D and 5E). We also detected slightly elevated neuronal Ca^2+^ PD in neuropil in higher frequency ranges (theta) during IS sleep (Figure 5E). Finally, we quantified the temporal relationship between astrocytic and neuronal Ca^2+^ signals within the same neuropil ROIs. We found rather low levels of synchronicity between the two cell types, and overall reduced correlation during sleep compared to wakefulness (Figure 5F). Taken together, astrocytic Ca^2+^ signaling in sleep differs from overall local neuronal activity.

### Astrocytic IP_3_-mediated Ca^2+^ signaling modulates sleep states, brain rhythms and sleep spindles

To identify the potential roles of astrocytic Ca^2+^ signals in sleep, we employed the *Itpr2*^−/−^ mouse model, in which astrocytic Ca^2+^ signaling is strongly attenuated but not abolished (Srinivasan et al., 2015). We compared sleep architecture and spectral ECoG properties in WT and *Itpr2*^−/−^ mice. We found that *Itpr2*^−/−^ mice exhibited more frequent NREM and IS bouts that were of shorter duration than in the WT mice (Figures 6A and 6B). Relative time spent in NREM and IS did not differ between the two genotypes (Figure 6C), indicating a more fragmented NREM and IS sleep in *Itpr2*^−/−^ mice. Conversely, frequency of REM bouts, REM bout duration and relative time spent in REM sleep were all similar in the two genotypes (Figures 6A–C), in accordance with a previous report (Cao et al., 2013). Next, we assessed the spectral ECoG properties of NREM, IS and REM sleep between the genotypes. We detected a 7% decrease in delta power and a 16% increase in sigma power in *Itpr2*^−/−^ mice during IS sleep, compared to WT mice (Figures 6D and 6F), suggesting that astrocytic IP_3_-mediated Ca^2+^ signaling may be involved in the regulation of NREM to REM sleep transition. Theta power was 11% higher in *Itpr2*^−/−^ mice during REM sleep than in WT mice (Figure 6E), proposing that IP_3_-mediated Ca^2+^ signals may dampen the theta rhythm. As sigma activity is indicative of sleep spindles – i.e. bursts of neuronal activity linked to memory consolidation (Diekelmann and Born, 2010) – we next evaluated the occurrence of sleep spindles in WT and *Itpr2*^−/−^ mice (Figure 6G). The frequency of sleep spindles in IS sleep was twice as high in *Itpr2*^−/−^ mice vs. WT mice (Figure 6H). These data support a role for astrocytic IP_3_-mediated Ca^2+^ signaling pathway in modulating sleep states, brain rhythms, and sleep spindle homeostasis.

**Figure 6.**
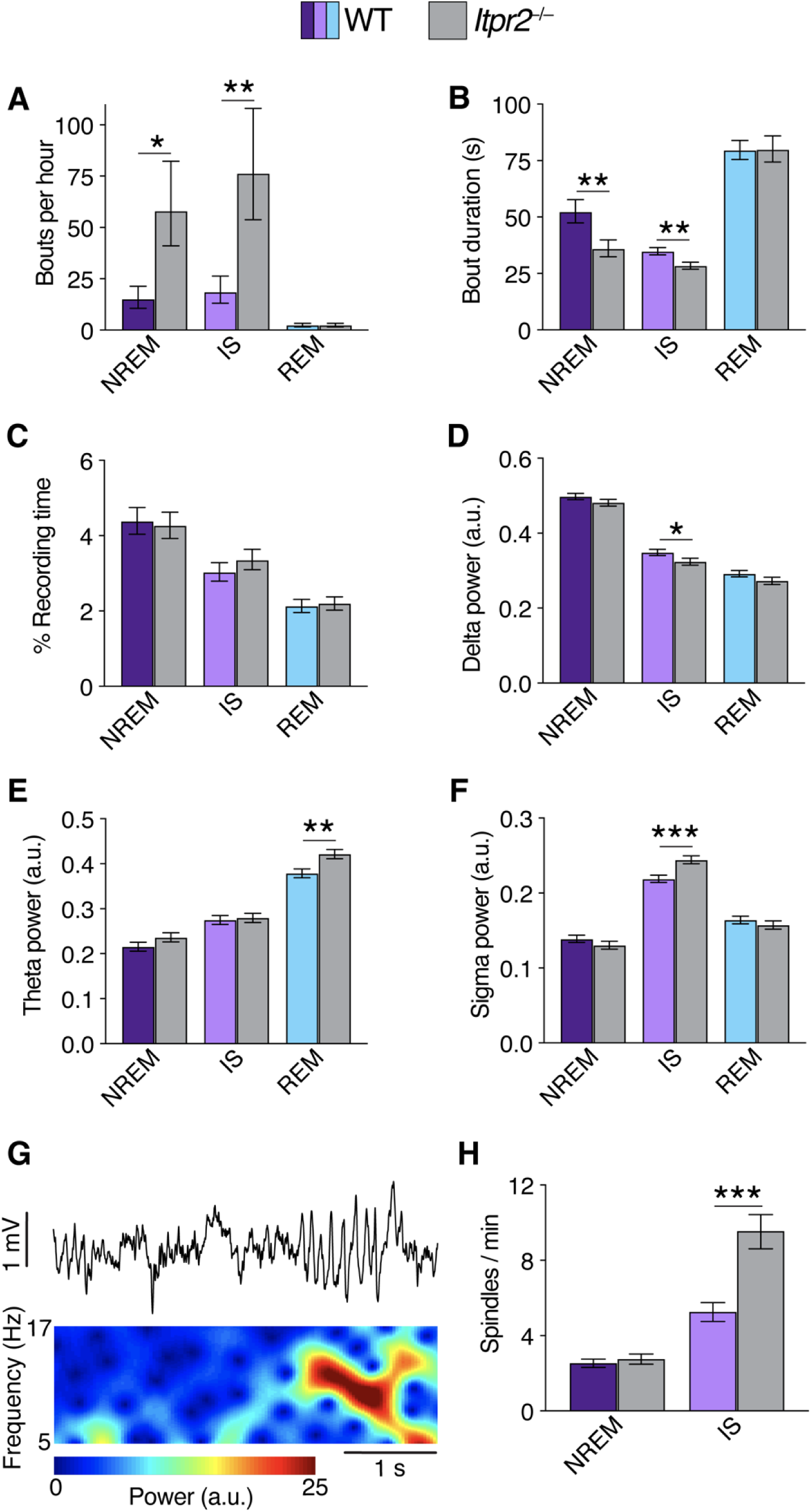
The astrocytic IP_3_-mediated Ca^2+^ signaling pathway modulates sleep architecture, brain rhythms and sleep spindles. (A to C) Mean number of bouts per hour (A), bout duration (B), and percentage recording time (C) of NREM, IS and REM states in WT and *Itpr2*^−/−^ mice. (D to F) Normalized mean power of delta (0.5–4 Hz) (D), theta (5–9 Hz) (E) and sigma (10–16 Hz) (F) frequency bands in NREM, IS and REM sleep in WT and *Itpr2*^−/−^ mice (normalized to 0.5–30 Hz total power). (G) Representative example of a sleep spindle in ECoG trace and power spectrogram. (H) Mean number of spindles per minute during NREM and IS sleep states in WT and *Itpr2*^−/−^ mice. All values represent estimates ± SEM, *p<0.05, **p<0.005, ***p<0.0005. For WT mice: n = 6 mice, 252 trials; for *Itpr2^−/−^* mice: n = 6 mice, 153 trials. For details on statistical analyses, see Methods.

### Ca^2+^ signaling during sleep and sleep-to-wake transitions is dependent on the IP_3_ pathway

In search for mechanisms explaining the IP_3_R2 mediated effects on sleep architecture and brain rhythm dynamics, we compared the characteristics of astrocytic Ca^2+^ signaling in *Itpr2*^−/−^ and WT mice (Figures 7 and S12). We found that *Itpr2*^−/−^ mice exhibited reduced astrocytic Ca^2+^ signaling in all states of wakefulness as measured by ROA frequency and active voxels (*x-y-t*) (Figure S12A and S12B), in agreement with previous reports (Srinivasan et al., 2015). Ca^2+^ events in *Itpr2*^−/−^ mice were of longer duration, but were smaller in area and volume (*x-y-t*) in all states of wakefulness compared to WT mice (Figure S12C–E). However, astrocytic Ca^2+^ activity measured by both the percentage of active voxels (*x-y-t*) and ROA frequency did not significantly differ between WT and *Itpr2^−/−^* mice during sleep (Figure 7B–C). Even so, astrocytic Ca^2+^ signals in *Itpr2*^−/−^ mice were of longer duration in all sleep states and of smaller spatial extent during NREM and IS sleep (Figures 7D and 7E). We did not detect differences in ROA volume in any sleep state compared to WT mice (Figure S13E).

**Figure 7.**
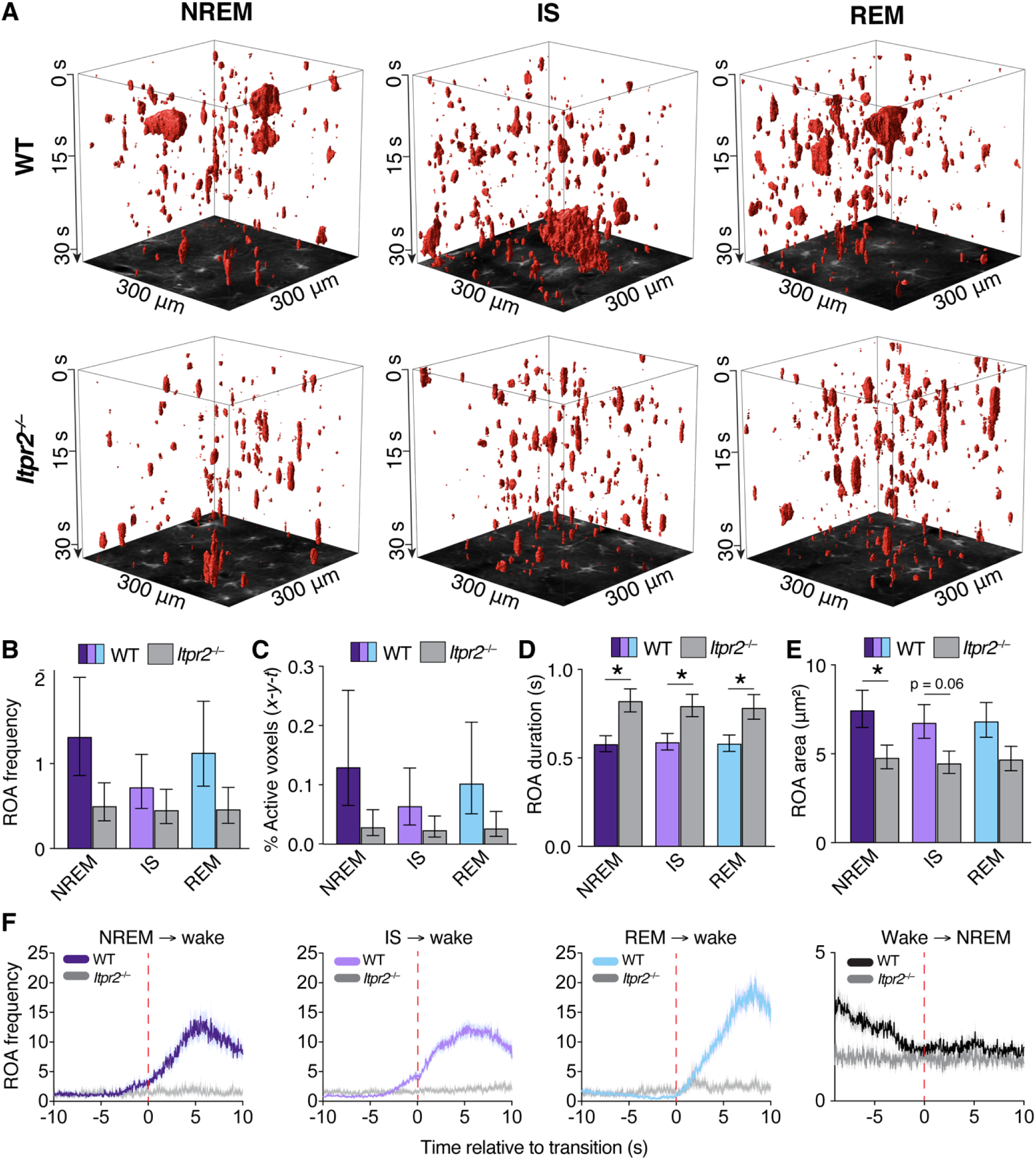
Astrocytic Ca^2+^ signaling during sleep is dependent on the IP_3_ pathway. (A) Representative *x-y-t* rendering of ROAs during NREM, IS and REM sleep in WT and *Itpr2*^−/−^ mice. (B to E) Mean ROA frequency expressed as number of ROAs per 100 μm^2^ per minute (B), the percentage of active voxels (*x-y-t*) (C), ROA duration (s) (D) and ROA area (μm^2^) (E) during NREM, IS and REM sleep in WT and *Itpr2*^−/−^ mice. Data represented as estimates ± SEM. For WT: n = 6 mice, 196 trials; for *Itpr2*^−/−^: n = 6 mice, 100 trials. (F) Mean time-course of astrocytic Ca^2+^ signals during transitions to wakefulness from NREM, IS and REM sleep, and from wakefulness to NREM sleep, WT and *Itpr2*^−/−^ mice. Data represented as mean ± SEM, n = 6 mice for WT, n = 6 mice for *Itpr2*^−/−^. *p<0.05, **p<0.005, ***p<0.0005. For details on statistical analyses, see Methods. See also Figures S12 and S13.

Interfering with astrocytic Ca^2+^ signaling also modulated the characteristics of neuronal Ca^2+^ signaling (Figures S13). Specifically, the amplitude of neuronal Ca^2+^ events was lower in *Itpr2*^−/−^ mice during all sleep states, but not during wakefulness (Figure S13C). The frequency and duration of neuronal Ca^2+^ events did not differ between the genotypes, neither during sleep nor wakefulness (Figures S13A and S13B). Moreover, neuronal Ca^2+^ oscillation power in the neuropil of *Itpr2*^−/−^ mice was lower in the delta range during SWS (NREM and IS), while higher in theta range in all sleep states (Figures S13D and S13E), supporting a role of astrocytic IP_3_ mediated Ca^2+^ signaling in promoting slow frequency rhythms and dampening high frequency rhythms during sleep (Foley et al., 2017; Poskanzer and Yuste, 2016).

As sleep-to-wake transitions were associated with prominent rises in Ca^2+^ signaling (Figure 4), we investigated Ca^2+^ signaling in *Itpr2*^−/−^ mice in relation to brain state transitions. *Itpr2*^−/−^ mice did not display increases in Ca^2+^ signaling in neither NREM / IS / REM sleep-to-wake transitions (Figure 7F). Moreover, falling asleep in *Itpr2*^−/−^ mice was not associated with a decrease in Ca^2+^ signaling, as seen in WT mice (Figure 7F). To summarize, astrocytic Ca^2+^ signaling during sleep and sleep-to-wake transitions is dependent on the IP_3_ pathway.

## DISCUSSION

Astrocytes are emerging as essential components of neural circuits that modulate awake behavior, particularly through Ca^2+^ signaling (Adamsky et al., 2018; Mu et al., 2019; Yu et al., 2018). In this work, we discovered that astrocytic Ca^2+^ signaling changes prominently across the sleep-wake cycle, being reduced during sleep while abruptly becoming elevated upon awakening, often before behavioral and neurophysiological signs of the sleep-to-wake transition. Genetic ablation of a key astrocytic Ca^2+^ signaling pathway altered sleep architecture and resulted in fragmentation of SWS, yet increased the frequency of sleep spindles. We propose that astrocytes are essential for normal sleep through mechanisms involving intracellular Ca^2+^ signals. The concept that a non-neuronal cell type is indispensable for appropriate sleep may guide future studies aimed at deciphering sleep regulatory mechanisms and identifying novel treatment strategies for sleep disorders.

Until now, astrocytic Ca^2+^ signals *in vivo* have only been studied in anesthetized and awake animals, with prominent differences in signaling patterns reported for the two brain states (Bindocci et al., 2017; Nimmerjahn et al., 2009; Thrane et al., 2012). The use of ultrasensitive genetically encoded Ca^2+^ indicators and two-photon microscopy has revealed that astrocytes in unanesthetized animals have a plethora of Ca^2+^ signals, predominantly occurring in their processes (Bindocci et al., 2017; Haustein et al., 2014; Srinivasan et al., 2015; Stobart et al., 2017). Since Ca^2+^ elevations within the complex morphology of astrocytes necessarily vary in location and extent over time, conventional analytical methods based on fixed ROIs are insufficient in capturing the signals. Here, we developed and employed a novel activity-based algorithm that enabled us to identify previously unreported characteristics of astrocytic Ca^2+^ signals during sleep and sleep-wake transitions. Our analytical tool was applied on images captured at a frame rate enabling correction of movement artifacts while maintaining a temporal resolution sufficient for detection of rapid (subsecond) astrocytic Ca^2+^ signals. Furthermore, the animals were continuously monitored with an IR camera, as well as ECoG and EMG measurements for reliable classification of behavior and brain states.

We separated wakefulness into three states – voluntary locomotion, spontaneous whisking and quiet wakefulness – and show that astrocytic Ca^2+^ signaling varies greatly across these states. Voluntary locomotion and spontaneous whisking were associated with a higher frequency of astrocytic Ca^2+^ events than quiet wakefulness. Locomotion, known to be associated with norepinephrine release and global activation of astrocytes (Paukert et al., 2014), triggered the highest Ca^2+^ signaling levels. Previously it was unclear whether physiological sensory input uncoupled from locomotion and startle elicits Ca^2+^ signaling in rodent cortical astrocytes. Notably, Ca^2+^ responses were only detected in barrel cortex astrocytes after high-velocity (90 Hz) whisker deflections or when the stimulus occurred together with volitional locomotion, but not during natural whisking alone (Stobart et al., 2018; Tran et al., 2018). Our demonstration of a robust astrocytic Ca^2+^ response to spontaneous whisking in resting animals likely relies on the use of our sensitive activity-based analytical tool. The finding is of particular importance since artificial whisker deflection and spontaneous whisking depend on distinct afferent inputs and signal processing mechanisms within the somatosensory cortex (Arabzadeh et al., 2005; Krupa et al., 2004). Our data also underscores the importance of careful behavioral monitoring when studying astrocytic Ca^2+^ signaling.

A surprising finding was the lack of increase in astrocyte Ca^2+^ signaling during REM sleep compared to NREM sleep. REM sleep is characterized by high cortical levels of extracellular acetylcholine compared to NREM sleep (Jasper and Tessier, 1971; Marrosu et al., 1995). Astrocytes have been shown to respond to acetylcholine with increases in Ca^2+^ signaling upon direct application or stimulation of cholinergic nuclei (Navarrete et al., 2012; Takata et al., 2011), we expected a strong increase in astrocytic Ca^2+^ activity during REM sleep. This was, however, not the case, suggesting that acetylcholine alone in physiological concentrations is not sufficient to trigger astrocyte Ca^2+^ elevations.

Ca^2+^ signaling was more frequent in astrocytic processes than in somata through all sleep-wake states, in line with previous findings *in vitro* and *in vivo* during anesthesia and awake behavior (Bindocci et al., 2017; Haustein et al., 2014). While cortical neurons show diversity in activity across sleep states (Niethard et al., 2016), we did not detect subpopulations of astrocytes or astrocytic microdomains maintaining high Ca^2+^ activity across multiple brain states. Instead, Ca^2+^ activity hot spots tended to shift location between states. To assess whether astrocytic Ca^2+^ signaling mirrored the activity of neighboring neurons, we simultaneously recorded neuronal Ca^2+^ signals, employing the human *synapsin1* promoter that ensured Ca^2+^ sensor expression in excitatory pyramidal cells as well as inhibitory interneurons. Overall, the frequency of Ca^2+^ signals in neuronal somata decreased from wake to sleep, but only REM sleep was accompanied by a reduced firing rate when compared to quiet wakefulness, similar to previous reports (Niethard et al., 2016). Furthermore, Ca^2+^ signaling in neuronal somata changed much less than the astrocytic activity. Finally, we did not detect a strong correlation between Ca^2+^ signals in astrocytes and neurons within the same FOV, and the correlation of the signals within neuropil was even lower during sleep than in wakefulness. Taken together, our findings suggest that astrocytic Ca^2+^ activity does not simply reflect the overall activity of local cortical neurons, but rather is regulated by intrinsic mechanisms or activity in specific neuronal circuits not identified with the present imaging strategy.

Our findings that astrocytic Ca^2+^ signaling displayed brain-state specific features and changed upon state transitions prompted us to test whether astrocytic Ca^2+^ signals modulate sleep. We applied the widely used *Itpr2^−/−^* knockout model, in which the removal of the IP_3_R2s suppresses spontaneous as well as evoked astrocytic Ca^2+^ signaling (Srinivasan et al., 2015). Notably, IP_3_R2 is expressed by glia, whereas IP_3_R1 and IP_3_R3 are predominantly neuronal (Sharp et al., 1999). Even though *Itpr2^−/−^* knockout more profoundly affected astrocytic Ca^2^ signaling during wakefulness than in sleep, it was associated with discernible changes in sleep characteristics and brain rhythms. Specifically, NREM and IS sleep bouts were more frequent but of shorter duration in mutant than in WT mice. At the same time *Itpr2^−/−^* mice had more sleep spindles. Our data suggests that attenuated or less widespread astrocytic Ca^2+^ signaling promotes SWS state shifts and spindle generation via effects on local cortical circuits. We did not study whether the manipulation of astrocytic Ca^2+^ signaling affected specific cortical neurons thought to regulate SWS and spindles (Niethard et al., 2016; Seibt et al., 2017). Furthermore, since the gene knockout was global, we cannot rule out that altered astrocytic Ca^2+^ signaling in distant (e.g. thalamic) regions affects neurons regulating SWS and spindles. Future studies should delineate the specific pathways and circuits responsible for astrocytic IP_3_ mediated modulation of sleep and resolve whether aberrant Ca^2+^ signaling in astrocytes underlies sleep disorders.

## Acknowledgements

We thank Xiaoyi Zhang for help with immunohistochemistry experiments. We are grateful to Johannes Helm and Arild Njå for technical assistance. We also thank Anna Chambers for reviewing the manuscript. We gratefully acknowledge the support by UNINETT Sigma2 AS, for making data storage available through NIRD, project NS9021K. This work was supported by the Research Council of Norway (grants #249988, #240476 and #262552), the South-Eastern Norway Regional Health Authority (grant #2016070), the European Union’s Seventh Framework Programme for research, technological development and demonstration under grant agreement no. 601055, The National Association of Public Health, The Olav Thon Foundation and the Letten Foundation.

## Author contributions

Conceptualization E.A.N., K.A.G.V.; Methodology L.B., R.E., K.A.G.V., E.A.N., W.T., R.S.; Software D.M.B., K.S.Å., R.E., K.H.P.; Formal analysis D.M.B., R.E., C.C., G.H.H., K.H.P., L.B.; Investigation L.B.; Resources W.T., E.A.N., R.S.; Writing – Original Draft L.B., R.E., E.A.N.; Writing – Review and Editing L.B., R.E., E.A.N.; Visualization L.B., R.E.; Supervision R.E., E.A.N., K.A.G.V.; Funding acquisition E.A.N..

## Declaration of Interests

The authors declare no competing interests

## Methods

### CONTACT FOR REAGENT AND RESOURCE SHARING

Further information and requests for resources and reagents should be directed to and will be fulfilled by the Lead Contact, Erlend Nagelhus.

### EXPERIMENTAL MODEL AND SUBJECT DETAILS

#### Animals

Male C57BL/6J (Janvier Labs), *Itpr2^−/−^* (*Itpr2*^tm1.1Chen^; MGI:3640970) (Li et al., 2005) and *Glt1*eGFP mice (Regan et al., 2007; Rosic et al., 2019) were housed on a 12-h light/dark cycle (lights on at 8 AM), 1–4 mice per cage. Each animal underwent surgery at the age of 8–10 weeks, followed by accommodation to being head-restrained and two-photon imaging sessions (2–3 times per week) for up to 2 months. Adequate measures were taken to minimize pain and discomfort. All procedures were approved by the Norwegian Food Safety Authority (project number: FOTS 11983).

### METHOD DETAILS

#### Cloning and virus production

Serotype 2/1 recombinant adeno-associated virus (rAAV) from plasmid constructs pAAV-*GFAP*-GCaMP6f (Chen et al., 2013; Enger et al., 2017) and pAAV*-hSYN*-jRGECO1a (Dana et al., 2016), were generated as described (Tang et al., 2009) and purified by AVB Sepharose affinity chromatography (Smith et al., 2009) following titration with real-time PCR (rAAV titers about 1.0–6.0 × 10^12^ viral genomes/mL, TaqMan Assay, Applied Biosystems Inc.). rAAV-*GFAP*-GCaMP6f and rAAV-*SYN*-jRGECO1a were mixed 1:1.

#### Surgical procedures and intrinsic imaging

On the first day, anesthesia was induced in a chamber containing 3% isoflurane in room air enriched with 50% pure oxygen and subsequently maintained by nose cone flowing 1.5–2% isoflurane. Buprenorphine 0.15 mg/kg was injected intraperitoneally and the mice were left for 10 min before surgery started. The field was sterilized, and local anesthesia was accomplished by injection of bupivacaine (5 mg/mL). Two silver wires (200 μm, non-insulated, GoodFellow) were inserted epidurally into 2 small burr holes overlying the right parietal hemisphere for electrocorticographic (ECoG) recordings, and two stainless steel wires (50 μm thickness, insulated, except 1 mm tip, GoodFellow) were implanted in the nuchal muscles for electromyographic (EMG) recordings. The skull over the left hemisphere was thinned for intrinsic signal imaging, a custom-made titanium head-bar was glued to the skull and the implant sealed with a dental cement cap. Mice received meloxicam (2 mg/kg) postoperatively for 2 days. After two days of recovery, representations of individual whiskers in the barrel cortex were mapped by intrinsic optical imaging through the thinned skull. For this procedure, mice were anesthetized with isoflurane (3% for induction, 1% for maintenance) and received chlorprothixene (1 mg/kg) amount of this substance intramuscularly. The brain region activated by single whisker deflection (10 Hz, 6 s) was identified by increased red light absorption (due to increased blood flow). After two days of recovery, chronic window implantation and virus injection was performed as described previously (Enger et al., 2017; Rosic et al., 2019). Mice received the same anesthesia and analgesia as on day one. A round craniotomy of 2.5 mm diameter was made over the barrel cortex using the intrinsic optical imaging map as a reference. The virus mixture (70 nL at 35 nL/min) was injected at two different locations in the barrel cortex at a depth of 400 μm relative to the brain surface. A window made of 2 circular coverslips of 2.5 and 3.5 mm was glued together by ultraviolet curing glue (Huber et al., 2012), was then centered in the craniotomy, so that the glass plug very slightly depressed the dura. The window was subsequently fastened by dental cement. Mice received meloxicam (2 mg/kg) postoperatively for 2 days.

#### Behavioral training

Mice were housed in an enriched environment with a freely spinning wheel in their home cages. One week before imaging, mice were habituated to be head-fixed on a freely spinning wheel under the two-photon microscope. Each mouse was trained head-fixed daily before the imaging for increasing durations ranging from 10 min on the first day to 70 min on the last. The mouse was removed from the head-restraint if it showed signs of stress over 20 min.

#### Two-photon imaging

Four weeks after the craniotomy surgery and virus transduction, mice were imaged under a laser scanning two-photon microscope (Ultima IV from Bruker/Prairie Technologies) as previously described (Enger et al., 2015, 2017; Rosic et al., 2019), with a Nikon 16 × 0.8 NA water-immersion objective (model CFI75 LWD 16XW). The fluorescence of GCaMP6f and jRGECO1a were excited at 999 nm with a Spectra-Physics InSight DeepSee laser, and emitted photons were detected with Peltier cooled photomultiplier tubes (model 7422PA-40 by Hamamatsu Photonics K.K.). All optical filters mentioned in the following description are by Chroma Technology Corporation: after having been reflected towards the detection unit by the main dichroic filter (type ZT473-488/594/NIRtpc), the signal light enters the system’s 4-channel detector house, at the entrance of which a type ZET473-488/594/NIRm filter is installed, shielding the photomultiplier tubes from rest reflective light of the laser beams. Inside the detector house, the light is split into 2 fractions separated at 560 nm wavelength by the main signal light dichroic filter (T560lpxr). The “green” light (GCaMP6f fluorescence) is further guided by a secondary dichroic beam splitter at 495 nm (T495lpxr) and filtered by a ET525/50m-2p band-pass filter, whereas the “red” light (jRGECO1a fluorescence) is similarly guided by a secondary beam splitter at 640 nm (T640lpxr) and subsequently filtered by a ET595/50m-2p band-pass filter. Images (512 x 512 pixels) were acquired at 30 Hz in layer 2/3 of barrel cortex. To assist sleep while head-fixed, the wheel was lifted higher towards the objective for more natural sleeping posture, and the movement of the wheel was locked once the mouse showed first signs of sleep. Mice that did not show any signs of sleep within the first 2 hours were removed from the microscope.

#### Behavior and electrophysiology recording

Data acquisition (two-photon imaging, ECoG, EMG and infrared-sensitive surveillance camera video) was synchronized by a custom-written LabVIEW software (National Instruments). ECoG and EMG signals were recorded using a Multiclamp 700B amplifier with headstage CV-7B, and digitized by Digidata 1440 (both Molecular Devices). ECoG and EMG data were analyzed using custom-written MATLAB scripts. Mouse behavior was recorded by an infrared-sensitive surveillance camera.

#### Sleep-wake state scoring

Sleep states were identified from filtered ECoG and EMG signals (ECoG 0.5–30 Hz, EMG 100– 1000 Hz) based on standard criteria for rodent sleep (Kohtoh et al., 2008; Kreuzer et al., 2015; Niethard et al., 2016; Seibt et al., 2017) (See Figure 1C–D). Non-rapid eye movement (NREM) sleep was defined as high amplitude delta ECoG activity (0.5–4 Hz) and low EMG activity; intermediate state (IS) was defined as an increase in theta (5–9 Hz) and sigma (9–16Hz) ECoG activity, and a concomitant decrease in delta ECoG activity; rapid eye movement (REM) sleep was defined as low amplitude theta ECoG activity with theta/delta ratio >0.5 and low EMG activity. Wakefulness states were identified using the infrared-sensitive surveillance camera video by drawing ROIs over the running wheel and mouse snout (Figure 1B). The signal in the wheel and snout ROIs was quantified by calculating the mean absolute pixel difference between consecutive frames in the respective ROIs. Voluntary locomotion was identified as signals above a threshold in the wheel ROI. Similarly, spontaneous whisking was defined in the snout ROI. Quiet wakefulness was defined as wakefulness with no signal above threshold in both ROIs. Only episodes of wakefulness of more than 10 s duration and sleep episodes of more than 30 s duration were analyzed.

#### Sleep-wake state transition scoring

Sleep-to-wake transitions were determined as previously described (Takahashi et al., 2006, 2009, 2010). For the transition from NREM and IS to wakefulness, onset of wakefulness was determined by the first sign of ECoG desynchronization (activation) (Figure 4B). During the transitions from REM to wakefulness, end of REM was identified by the interruption of sustained theta waves and the onset of desynchronized ECoG (Figure 4B). Quiet wakefulness to locomotion was defined using the surveillance video (see above).

#### Immunohistochemistry

Mice were deeply anesthetized with isoflurane and intracardially perfused with phosphate buffered saline (PBS; 137 mM NaCl, 2.7 mM KCl, 4.3 mM Na_2_HPO_4_⋅2H_2_O, 1.4 mM KH_2_PO_4_, pH 7.4, all from Sigma-Aldrich) and 4% paraformaldehyde (PFA, Merck) prior to decapitation. Brains were removed and fixed in ice-cold 4% PFA/PBS for overnight. Serial 70 μm sections were obtained on a vibratome (Leica). Free-floating sections were washed in PBS, incubated for 1 h in blocking solution (4% normal goat serum (NGS) and 0.3% Triton X-100 in PBS), washed with PBS, incubated overnight at room temperature with primary antibodies in PBS-Triton with 1% NGS (Primary antibodies: polyclonal chicken anti-GFP (1:3000, Abcam), rabbit anti-GFP (1:4000, Abcam), mouse anti-NeuN (1:1000, Merck), mouse anti-GFAP (1:1000, Merck), rabbit anti-Iba1 (1:1000, Wako), washed in PBS, transferred into secondary antibody solution for 45 min (Secondary antibodies: Cy5-coupled anti-rabbit, Cy5-coupled anti-mouse, FITC-coupled anti-rabbit and FITC-coupled anti-chicken (1:200, Jackson ImmunoResearch Labs), washed with PBS and mounted on slides with Quick-hardening Mounting Medium (Sigma Aldrich). Confocal images were acquired on a Zeiss LSM710/Elyra S1 confocal laser scanning microscope equipped with a 4x air objective or 20x water objective. Image analysis was done with ImageJ (v10.2, NIH).

### QUANTIFICATION AND STATISTICAL ANALYSIS

#### Ca^2+^ signal analyses

Imaging data were corrected for motion artifacts using the NoRMCorre movement correction software (Pnevmatikakis and Giovannucci, 2017). Images were analyzed by a custom-made image-processing toolbox in MATLAB (Mathworks).

#### Region-of-activity (ROA) algorithm

For the automatic ROA analysis, data underwent the following steps (see Figure S2 for references): (A) *Preprocessing the imaging data.* Imaging data was corrected for movement artifacts and slightly smoothed in the spatial domain (gaussian smoothing, σ = 2 pixels) producing the time series *F*. (B) *Calculating ΔF/F_0_ time series.* A baseline image (*F_0_*) was calculated by first smoothing the pre-processed time series (*F*) in time (moving average filter, width 1.0 second), resulting in a lowpass filtered time series (*F_LP_*), and then subsequently calculating the mode of each of the pixels over time. The pre-processed time series (*F*) was then subtracted and divided by the baseline image (ΔF/F_0_ = (*F* - *F_0_*) / *F_0_*), resulting in a ΔF/F_0_ time series (*S*). (C) *Calculating a noise-based activity threshold and thresholding the data.* A moving average filter (width, *w* = 1.0 second) was then applied to the ΔF/F_0_ time series to create a smoothed, lowpass filtered time series (*S_LP_*). A highpass filtered time series (*S_HP_*) was then created by subtracting the smoothed time series (*S_LP_*) from the original ΔF/F_0_ time series (*S*). The highpass filtering of S was done to estimate noise in our time series. In an attempt to estimate the variance of the noise in *S* we assumed that it could be approximated by variance of *S_HP_*. As *S_LP_* is a moving average filtered version of *S* we would then expect that the standard deviation of the noise in *S_LP_* is a factor of 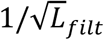 lower than *S*, where *L_filt_* is the length (number of frames) of the moving average filter. A standard deviation image (*σ*) (the noise approximation of *S_LP_*) was created by calculating the standard deviation of *S_HP_* and diving by 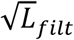. Artifacts from fluorescence drifts (Niethard et al., 2016) were removed by subtracting a 10 second moving average of *S_LP_* from *S_LP,_*, producing a final, bandpass filtered time series *S_BP_*. Voxels in *S_BP_* were then thresholded by the corresponding pixels in the σ image and multiplied by a common factor *k* (*k* = 5 was used in the analysis), resulting in a 3D matrix of active or inactive voxels. (D) *Detect connected components and extract descriptive properties of the events.* Adjacent, active voxels in space and time were assigned to single ROAs. As vessel walls move within the field-of-view with vascular dilation and constriction, the regions over and immediately surrounding blood vessels were prone to artifacts and manually masked out before connecting voxels. For each ROA the starting time, maximal spatial extent (µm²), volume (µm² · s) and duration (s) was recorded.

ROA frequency was calculated by counting all ROAs with starting times within a given frame and subsequently dividing by the sampled area (total field-of-view minus areas over vessels and their immediate surrounding) and the time per frame. The percentage of active voxels in a particular state episode was calculated by dividing the number of active voxels by the total number of voxels in that episode, while excluding the ignored areas. 3D renderings of ROA activity (Figure 2) were made by outlining the ROAs and plotting with the MATLAB function *patch()*. The time resolution was decimated for improved performance and visual representation.

ROA heatmaps were created by calculating the percentage of active voxels along the temporal dimension. Values lower than 0.3 % were left transparent while showing the average fluorescence of the astrocyte channel underneath (Figure 3). To assess the persistence of hotspots in the heatmaps the 5% most active pixels were defined, and the overlap between different behavioral states were calculated. One heatmap was calculated per behavioral state for each unique FOV (the same FOV could span multiple recordings). For each heatmap a binary image was created with the 5% most active pixels marked as true. The overlap between the binary images were then calculated separately between the wake states (locomotion / whisking / quiet), between the sleep states (NREM / IS / REM), between wake and sleep states (whisking / quiet / NREM / IS / REM) and between the merged wake and merged sleep states (wake / sleep). The overlap was defined as the area of the intersection divided by the area of the union.

#### Hand-drawn ROI analysis

Neuropil ROIs were marked as circles of 5 μm radius at least 5 μm away from the perimeter of astrocyte somata in areas without large visible cell processes. Astrocyte and neuron soma ROIs were manually drawn. Mean fluorescence traces (F) were calculated by averaging the fluorescence intensity inside the respective ROIs. The baseline (F_0_) of the traces was defined as the mode of the traces rounded to the nearest integer (the imaging data was in the format of unsigned 12-bit integers). Relative fluorescence was defined with the formula ΔF/F_0_ = (F-F_0_)/F_0_. For neuron soma signals, ΔF/F_0_ was calculated slightly differently to prevent ‘contamination’ from surrounding neuropil signals: We subtracted mean fluorescence in a peri-somatic ROI from the soma signal. Peri-somatic fluorescence traces were calculated from 5 μm wide regions around the neuron soma ROIs (‘doughnut’) and fitted to the neuron soma traces using least squares (F_doughnut_). The relative fluorescence change for neuron soma was then defined as ΔF/F_0_ = (F-F_doughnut_)/F_0_.

The relative change in fluorescence (ΔF/F_0_) from each ROI was filtered using a gaussian filter (σ = 0.25 s for GCaMP6f, σ = 0.1 s for jRGECO1a). To define thresholds for event detection, traces of the noise was approximated by subtracting the smoothed trace from the raw ΔF/F_0_ trace. In addition, a heavily smoothed fluorescent trace (gaussian filter, σ = 120 s) was subtracted from the lightly smoothed ΔF/F_0_ to remove drift. Increases in ΔF/F_0_ were identified as Ca^2+^ events if the increases were larger than 2.5 times of the GCaMP6f noise trace and 2 times the standard deviation of the *SYN*-jRGECO1a noise trace. The duration of the signal was defined as the earlier and later time points where the ΔF/F_0_ crossed 0 and the amplitude was the peak of the ΔF/F_0_ within the duration.

#### Ca^2+^ analyses in relation to brain state transitions

Traces of ROA frequency were smoothed to reduce the effect of noise crossing the threshold. As we were concerned that a conventional gaussian kernel would skew the onset estimates to earlier values we instead used a kernel on the form *t* ⋅ *exp*(−*t* / *τ*) (*τ* = 0.25 s) that rises quickly, and tapers off with an exponential decay. The traces in a window (-15 to 15 seconds) around every transition were collected to estimate the peak ROA frequency and the onset of Ca^2+^ activity. The peak ROA frequencies were estimated from the maximum value of the ROA frequency traces. ROA frequency was normalized via z-score transformation, which were the mean and standard deviation was calculated from the first 10 seconds (-15 to -5 seconds from the transition). The onset of Ca^2+^ activity was then measured at the first point the traces crossed a threshold of 2.5 standard deviations.

#### Neuropil Ca^2+^ power analysis

ΔF/F_0_ traces from *SYN*-jRGECO1a were extracted from circular 5 μm radius neuropil ROIs (see above, hand-drawn ROI analysis) for each 10 min trial using *mode* to calculate F_0_. Neuropil signal was calculated as the average of all neuropil ROIs in the recordings. The power of delta (0.5–4 Hz), theta (5–9 Hz) and sigma (10–15Hz) frequency bands of each episode was then calculated using the built-in MATLAB function *bandpower()* and power spectrum using *pspectrum()*.

#### Ca^2+^ signal correlation analysis

For this analysis, neuropil ROIs were applied to the neuron and astrocyte channel to create pairs of ΔF/F_0_ signals. F_0_ was estimated by finding the *mode* of the fluorescent traces (using the builtin MATLAB function *mode()* on a signal rounded to 1 decimal place). The mean correlation coefficient was then calculated from the pairs of traces for each episode. To investigate correlations in frequency bands, the instantaneous power in delta, theta and sigma range was estimated. This was done by bandpass filtering the signals (zero-phase 4^th^ order Butterworth filter) in delta, theta and sigma range followed by calculating the square absolute value of the Hilbert transform. The resulting traces were used to calculate the mean correlation coefficient in each episode.

For the correlation between astrocytic events in the neuropil and the neuropil Ca^2+^ signal (Figure 5C) one pair of traces were created for each recording. The ROAs inside all neuropil ROIs were concatenated into a single ROA frequency trace (which was calculated by counting the number of events within all neuropil ROIs in a given frame and subsequent division by time step per frame and the area covered by the neuropil ROIs). For the neuronal Ca^2+^ signal, the neuropil ROIs (i.e. same ROIs used for astrocyte signals) were used to extract one ΔF/F_0_ trace from the neuron channel, where the baseline was estimated by the *mode* of the trace. Both neuronal and astrocytic traces were subsequently smoothed with a gaussian filter (σ = 0.5 s). Finally, the correlation coefficient was calculated for each episode.

The correlation coefficients were determined by the maximum Pearson cross-correlation between the traces. The mean lag in all comparisons were less than a single frame.

#### ECoG and sleep architecture analysis

ECoG power spectrograms of every sleep-wake state were calculated using the MATLAB function *pspectrum()* with default parameters from 0 to 256 Hz. To compare ECoG power across genotypes and different mice, raw data were normalized to the average total power in the 0.5–30 Hz frequency range during NREM sleep per mouse and average power was calculated using the MATLAB *bandpower()* function. To detect sleep spindles the ECoG signal was first normalized as above, followed by bandpass filtering in the frequency range 10–16 Hz (2^nd^ order zero-phase Butterworth filter). The analytic signal was then found by using the Hilbert transform, as a measure of the instantaneous ‘power’ of the bandpass filtered signal, and subsequently smoothed by a gaussian filter with a sigma of 0.2 s. Peaks in the smoothed data (found by the MATLAB function *findpeaks()*) were treated as putative sleep spindles. Peaks with a width at threshold of < 0.5 s or > 5 s were discarded. Sleep bout frequency, duration and relative time spent in the different sleep states were estimated by MATLAB *fitglm()* mixed effects model on log-transformed response variables with genotype and state and their interaction as fixed effects and mouse identity (for frequency and relative time) or state grouped by mouse identity (for duration) as random effects. Goodness of fit was evaluated by residual plots and the MATLAB *compare()* function.

#### Statistical analyses

Statistical analyses were conducted in R (version 3.6.0). This section describes the analysis of the following variables: percentage of active voxels (*x-y-t*), ROA frequency, ROA duration, ROA area, ROA volume, ROI frequency (Ca^2+^ events per minute per ROI), ROI duration, ROI amplitude and astrocyte-neuron correlation in neuropil ROIs (Figures 5G and 7L–N). We used regression models of the mixed effect type. As fixed effects, we included brain state (either the binary overall sleep / overall wake or the sleep-wake sub-states REM/IS/NREM/quiet wakefulness/whisking/locomotion), genotype (WT / *Itpr*2^−/−^), and the interaction between these two categorical predictors. In relevant cases, we also included cell type and subcellular compartment (astrocytic somata / astrocytic processes / neuron somata) as a fixed effect and the three-way interaction between state, genotype and cell type (and all underlying two-way interactions). For the ROA frequency and the percentage of active pixels we adjusted for differences in zoom factor by including the area covered by a pixel as a fixed predictor. We included individual mice as a random effect influencing the fixed effect of state (the binary sleep/wake version). For most responses, the fitted models indicated considerable heterogeneity between the mice.

For all variables except ROI frequency, we assumed ordinary linear mixed effect models for the log-transformed responses. The models were fitted using the nlme package (Pinheiro et al., 2019). The adequacy of model assumptions was investigated by various residual plots (Pinheiro and Bates, 2006). In the cases were the residual plots indicated deviations from the assumption of constant residual variance, we extended the model by allowing the residual variance to vary as a function of genotype and state. For the ROI frequency analysis, we fitted separate models for astrocytic somata, astrocytic processes and neuronal somata. For astrocytic somata and astrocytic processes, there was a non-negligible probability of observing zero events. Therefore, we conducted these analyses using a two-part mixed effects model, fitted by the GLMMadaptive package (Rizopoulos, 2019). One part models the probability of an observation being equal to zero, the other part models the non-zero observations. In our case, we chose a log-normal model for the non-zero observations and used the same fixed and random effects as described above. For the zero-part of the model we used the same fixed effects (state and genotype), but without interaction. The reported results from this analysis concern the non-zero part of the model. For each of the responses, we have reported the estimated response in different combinations of state and genotype. These effect estimates are linear combinations of the fixed effect estimates from the fitted models. For the responses that were analysed on the log scale the effect estimates have been transformed back to the scale of the observations. Approximate standard errors were found in the usual fashion from the inverse observed Fisher information matrix. The standard errors for the effect estimates were computed on the log scale and then transformed to the scale of observations. The error bars in the figures are therefore asymmetric. The reported p-values are based on the t-distribution, with degrees of freedom as provided from the nlme package. This is a common choice, but note that these tests only hold approximately for linear mixed effects models. No corrections for multiple comparisons were applied.

The analysis of neuropil power density was done in MATLAB using a linear mixed effect model with log-transformed responses with each sub-state as fixed effects. We included individual mice as a random effect influencing the fixed effect of state.

For the Ca^2+^ analyses in relation to brain state transitions, the peak ROA frequency and the onset of Ca^2+^ activity was estimated using linear regression models with peak frequency and onset as responses.

### DATA AND SOFTWARE AVAILABILITY

The code supporting the current study has not been deposited in a public repository because the code is part of a larger analysis suite that is under development, but the relevant code is available from the corresponding author on request.

### KEY RESOURCES TABLE

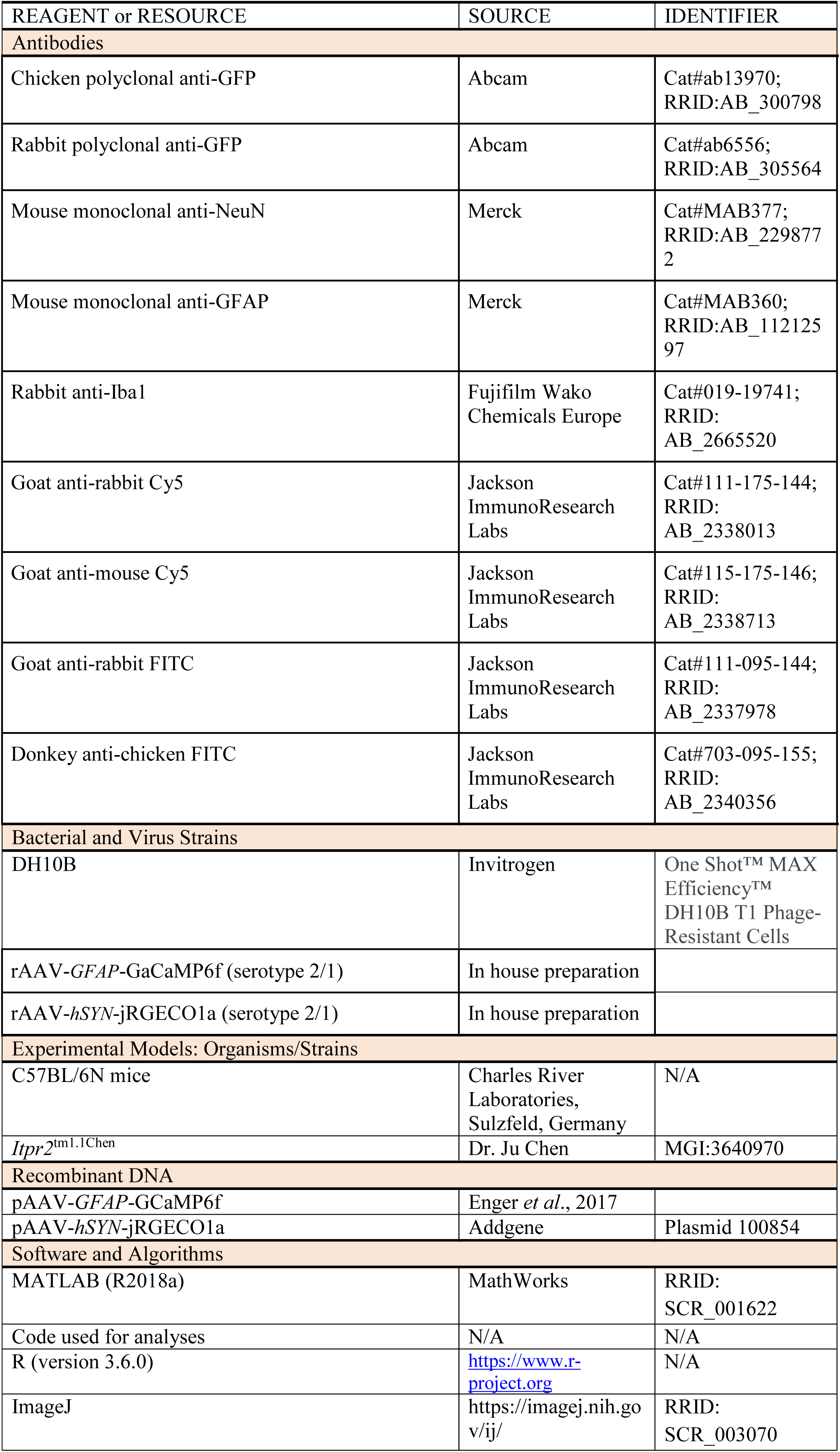

