## Supplementary Information for "Ca^2+^ signaling in astrocytes is sleep-wake state specific and modulates sleep"

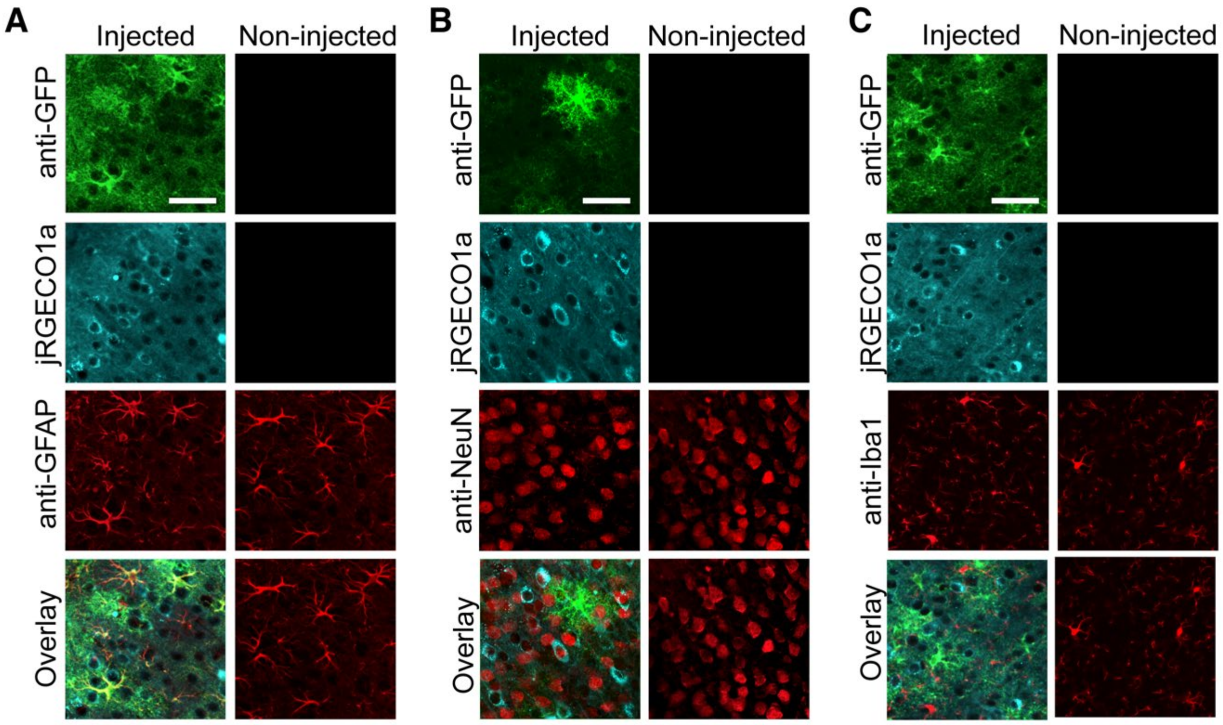


**Figure S1. Selectivity of the expression of *GFAP*-GCaMP6f in astrocytes and *SYN*-jRGECO1a in neurons.** (A) Immunolabeling of green fluorescent protein (anti-GFP, green) and glial fibrillary acidic protein (anti-GFAP, red), as well as jRGECO1a fluorescence (turquoise) demonstrate the selectivity of astrocytic and neuronal expression of the respective fluorescent indicators, and no astrogliosis, as compared to the contralateral non-injected hemisphere. Scale bar 50 µm. (B) Immunohistochemistry with anti-GFP and anti-NeuN, and fluorescence signal of jRGECO1a show that neurons, and not astrocytes, in the virus injection site were labeled with jRGECO1a. Scale = 50 µm. (C) Immunolabeling of GFP (anti-GFP, green) and the microglial marker Iba1 (anti-Iba1, red), as well as jRGECO1a fluorescence, demonstrate that injection of GCaMP6f and jRGECO1a did not induce microglial activation. Scale bars 50 µm.

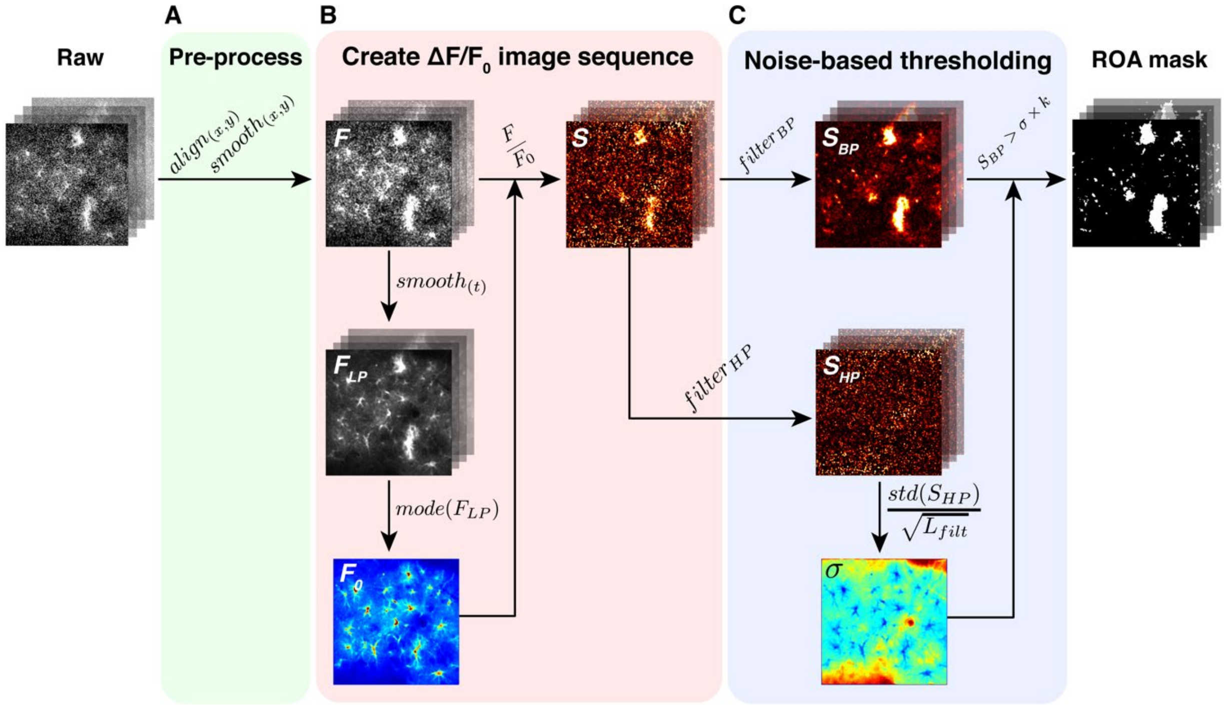


**Figure S2. Extracting regions-of-activity (ROAs) of astrocyte $Ca^{2+}$ signals from *GFAP*-GCaMP6f fluorescence images.** (A) The raw time series is pre-processed via image alignment and short spatial filtering ($\sigma = 2$ pixels). (B) A baseline image ($F_0$) is calculated by first smoothing the time series (moving average width = 1.0 s), resulting in a lowpass filtered time series ($F_{LP}$), and then subsequently calculating the mode of the pixels. The pre-processed time series F, is then divided by the $F_0$ image, resulting in a $\Delta F/F_0$ time series. (C) The $\Delta F/F_0$ time series is bandpass filtered in time to remove noise and slow drifts in baseline ($S_{BP}$). A highpass-filtered time series ($S_{HP}$) is used to estimate the noise and to calculate a standard deviation image ($\sigma$). Each pixel in the bandpass filtered time series ($S_{BP}$) is then thresholded by the corresponding pixel in the $\sigma$ image multiplied by a factor *k*.

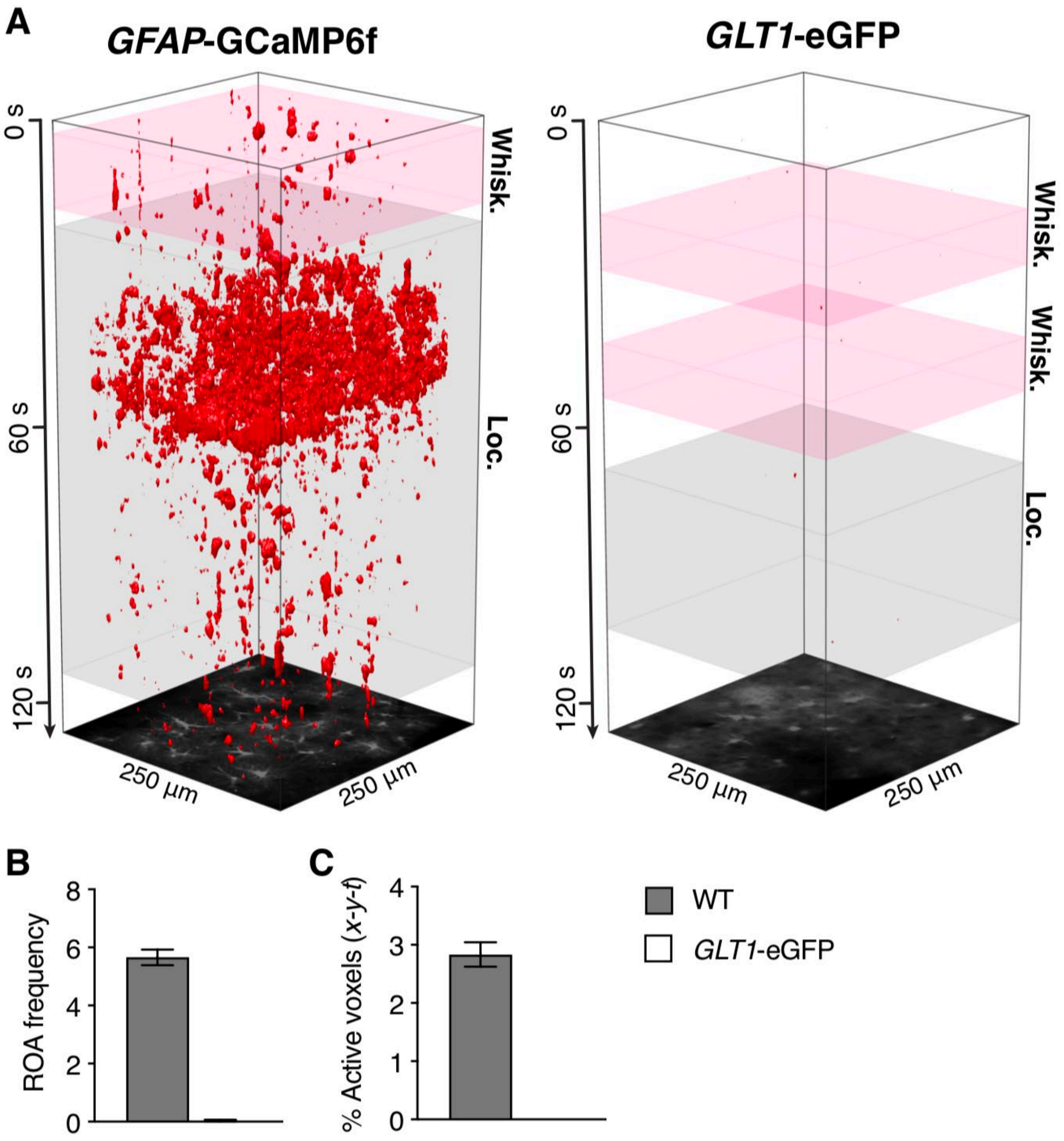


**Figure S3. Specificity of the ROA algorithm.** (A) Representative *x-y-t* rendering of ROAs detected in *GFAP*-GCaMP6f expressing astrocytes (A) and in eGFP expressing astrocytes (B) in wakefulness during locomotion (Loc.) and whisking (Whisk.). (C–D) ROA frequency (C) expressed as number of ROAs per 100 μm$^2$ per minute and the percentage of active voxels (*x-y-t*) (D) in the two genotypes. All bar graphs are represented as mean ± SEM, n = 6 mice, 82 trials for WT; n = 3 mice, 12 trials for *GLT1*-eGFP.

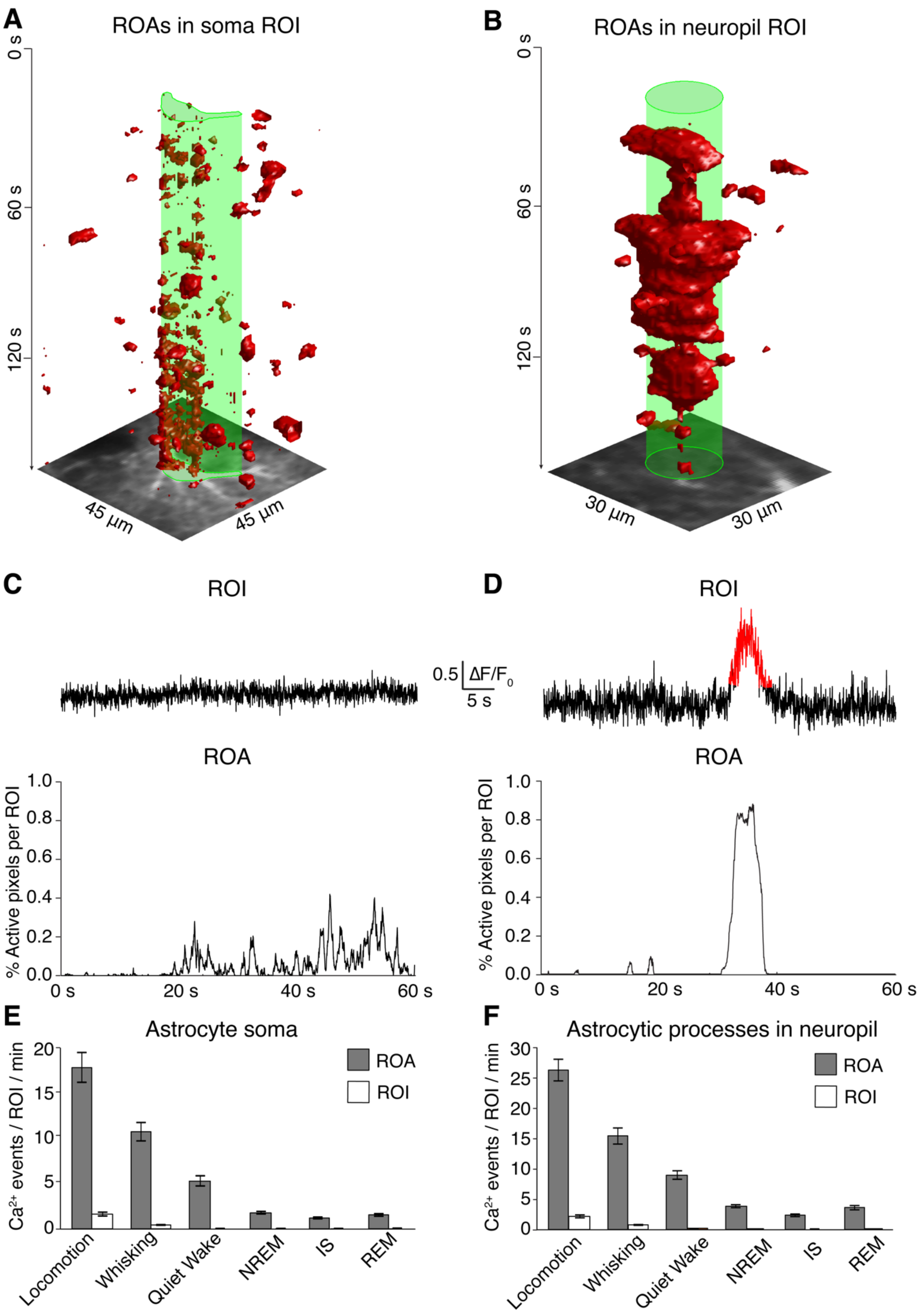

A
ROAs in soma ROI
0 s
60 s
120 s
45 µm
45 µm
B
ROAs in neuropil ROI
0 s
60 s
120 s
30 µm
30 µm
C
ROI
0.5 ΔF/F0
5 s
ROA
% Active pixels per ROI
1.0
0.8
0.6
0.4
0.2
0.0
0 s
20 s
40 s
60 s
D
ROI
ROA
% Active pixels per ROI
1.0
0.8
0.6
0.4
0.2
0.0
0 s
20 s
40 s
60 s
E
Astrocyte soma
Ca2+ events / ROI / min
20
15
10
5
0
ROA
ROI
Locomotion
Whisking
Quiet Wake
NREM
IS
REM
F
Astrocytic processes in neuropil
Ca2+ events / ROI / min
30
25
20
15
10
5
0
ROA
ROI
Locomotion
Whisking
Quiet Wake
NREM
IS
REM

**Figure S4**. **Comparison of region-of-interest (ROI) vs. ROA analyses.** (A and B) *x-y-t* rendering of ROAs in a ROI over astrocyte soma (A) or neuropil (containing astrocytic processes) (B). (C) Top: $\Delta F/F_0$ trace from the astrocyte soma ROI outlined in green. Bottom: percentage active pixels per ROI over time detected with the ROA algorithm. (D) Same as (C), but in neuropil ROI. (E) $Ca^{2+}$ event frequency per astrocyte soma ROI detected by standard ROI analysis compared to the new ROA analysis. (F) Same as (E), but in neuropil ROI. Data represented as mean ± SEM n = 6 mice, 243 trials.

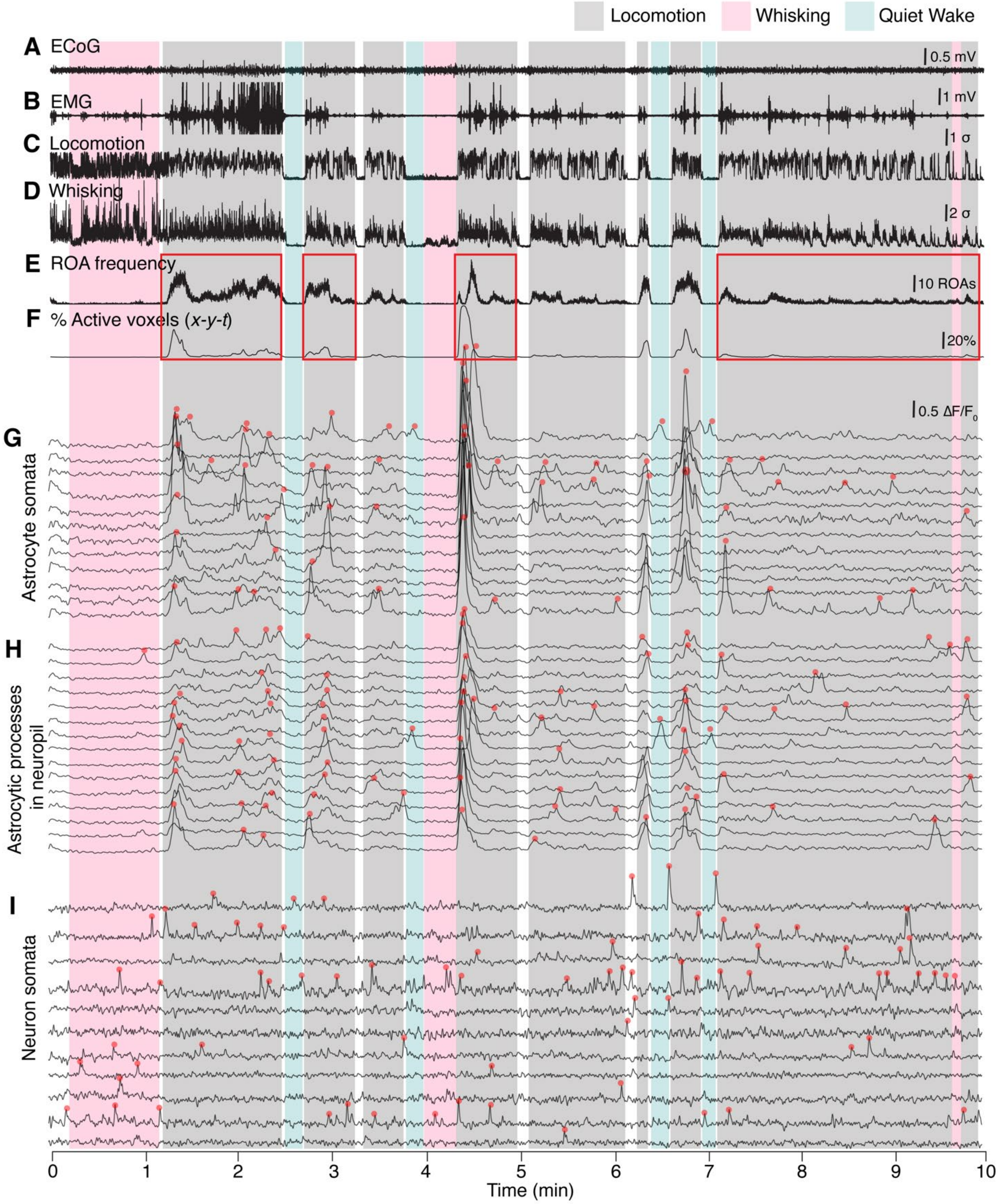


**Figure S5. A representative wakefulness trial.** (A) Bandpass filtered ECoG signal (0.5–30 Hz). (B) EMG signal. (C) Locomotion signal from a ROI placed on the wheel in the infrared-sensitive surveillance camera video. See also Figure 1B. (D) Whisker movement detected infrared-sensitive surveillance video. See also Figure 1B. (E) ROA frequency. (F) Percentage of active voxels (*x*-*y*-*t*). (G) $\Delta F/F_0$ traces from ROIs drawn over astrocyte somata. See also Figure S8. (H) $\Delta F/F_0$ traces for astrocytic $Ca^{2+}$ signals in neuropil ROIs, including astrocytic processes. See also Figure S8. (I) $\Delta F/F_0$ traces for ROIs drawn over neuron somata. See also Figure 5. Red squares illustrate higher astrocytic $Ca^{2+}$ activity related to the beginning of locomotion.

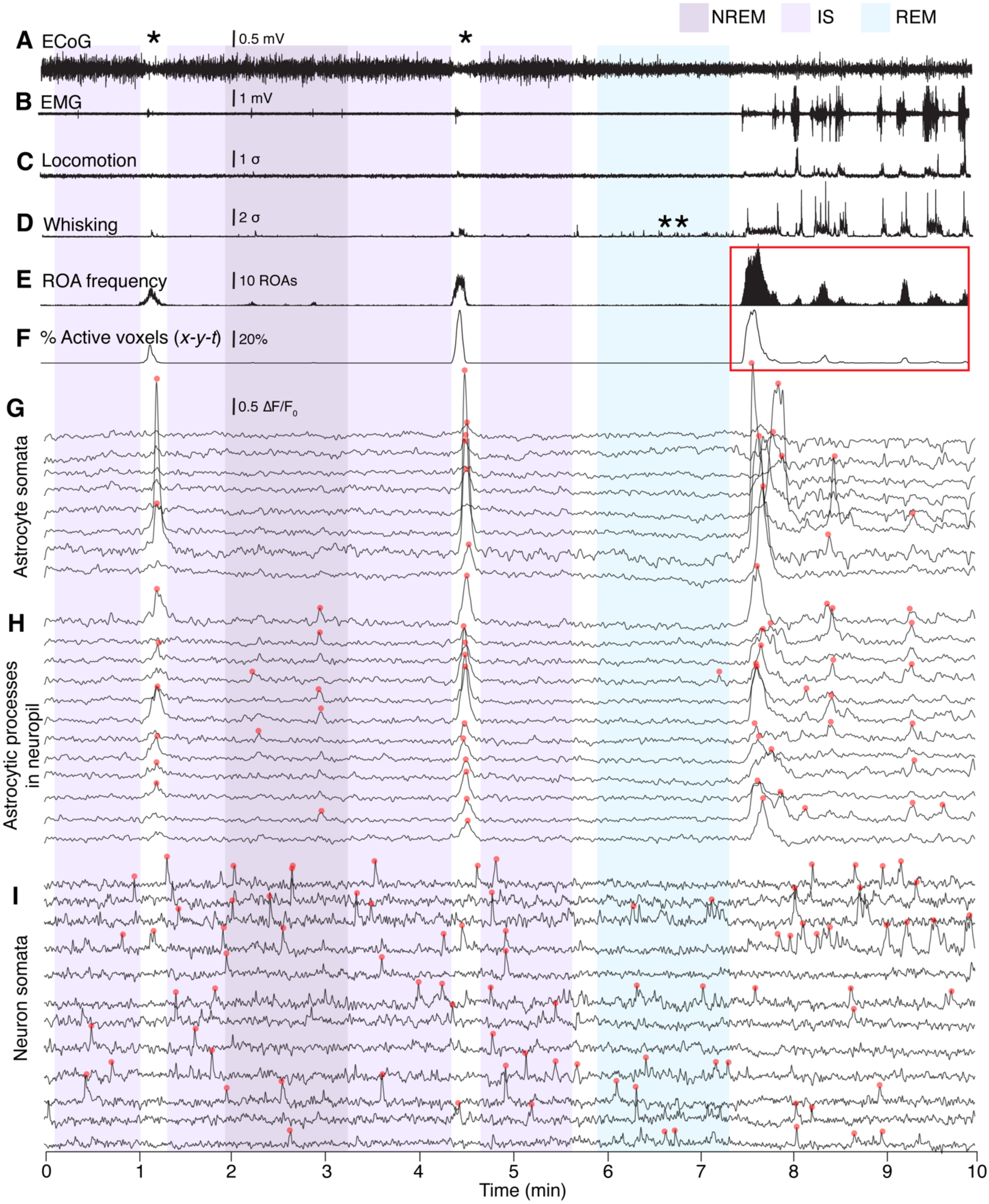


**Figure S6. A representative sleep trial.** (A) Bandpass filtered ECoG signal (0.5–30 Hz). Note an increase in ECoG amplitude during NREM and IS sleep. (B) EMG trace. Note absence of EMG activation during REM sleep. (C) Locomotion signal from a ROI placed on the wheel in the infrared-sensitive surveillance camera video. See also Figure 1B. (D) Whisker movement detected infrared-sensitive surveillance video. See also Figure 1B. **Note an increase in whisker signal during REM sleep without EMG activation. (E) ROA frequency. (F) Percentage of active voxels (*x*-*y*-*t*). (G) $\Delta F/F_0$ traces from ROIs drawn over astrocyte somata. See also Figure S8. (H) $\Delta F/F_0$ traces for astrocytic $Ca^{2+}$ signals in neuropil ROIs, including astrocytic processes. See also Figure S8. (I) $\Delta F/F_0$ traces for ROIs drawn over neuron somata. See also Figure 5. Red squares illustrate increased astrocytic $Ca^{2+}$ activity during an awakening. A single asterisk (*) indicates microarousals.

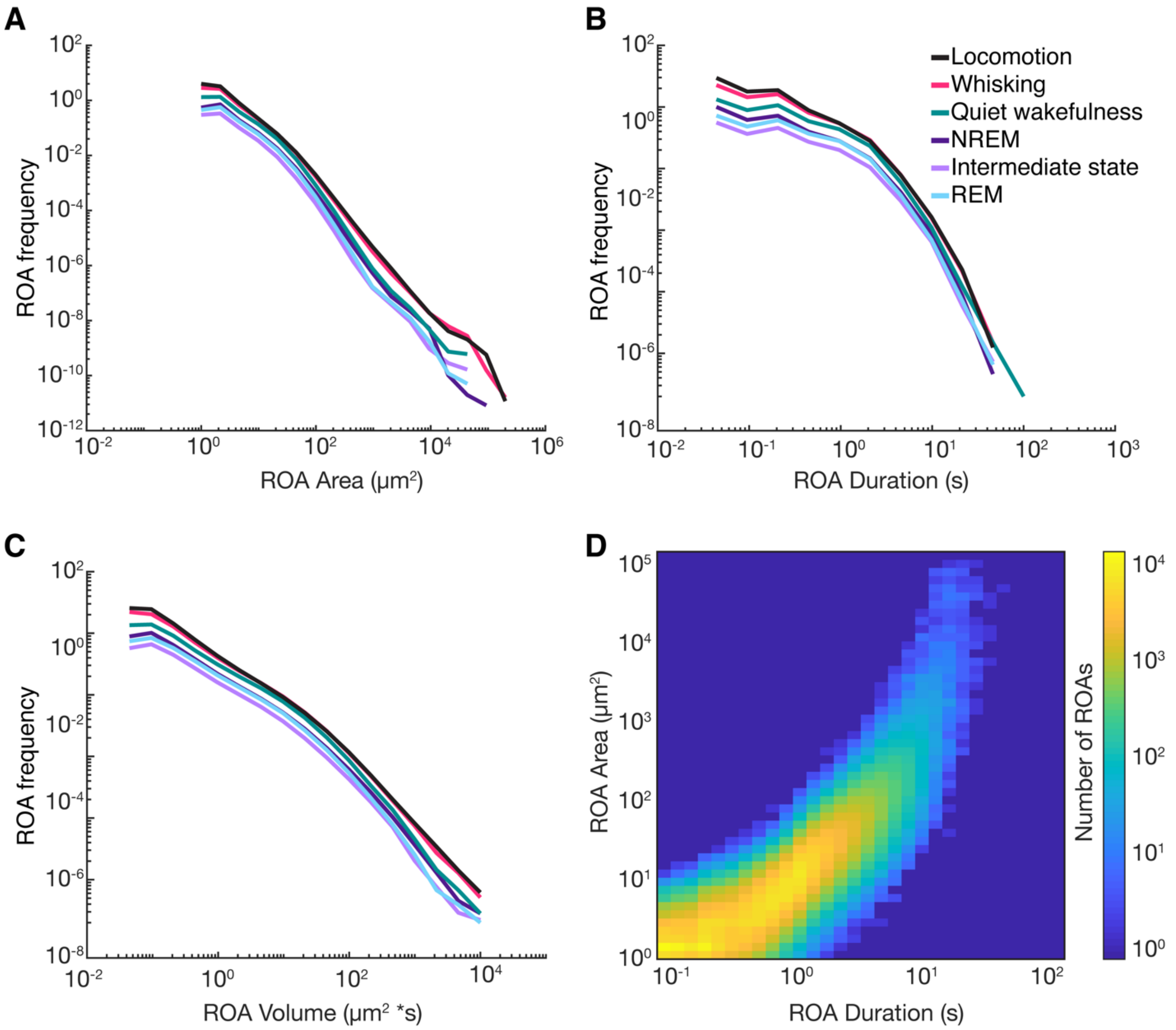


**Figure S7. Spatiotemporal characteristics of ROAs.** (A to C) The relationship between ROA frequency (y-axis) and ROA area (A), ROA duration (B) and ROA 3D (*x*-*y*-*t*) volume (C). (D) Bivariate histogram of ROA area and duration including ROAs from all sleep-wake states. n = 6 mice, 278 trials, 2 318 567 ROAs.

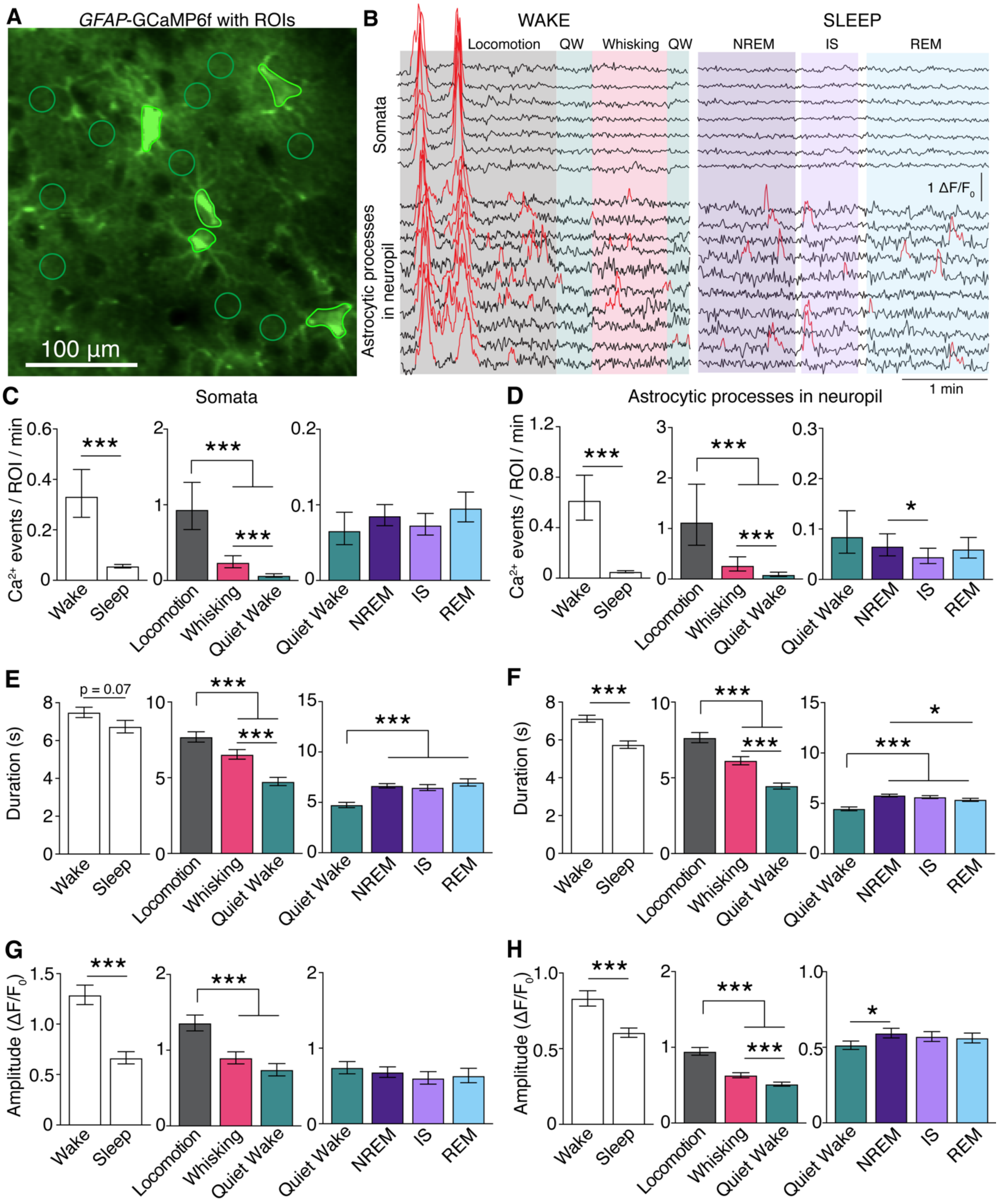


**Figure S8. Astrocytic $Ca^{2+}$ events across sleep-wake states from hand-drawn ROIs.** (A) Representative average image of *GFAP*-GCaMP6f fluorescence in astrocytes, and ROIs over astrocyte somata and neuropil. (B) Example $\Delta F/F_0$ $Ca^{2+}$ traces from astrocyte somata and neuropil ROIs across sleep-wake states. Detected $Ca^{2+}$ peaks are indicated in red. Traces smoothed by a 30-frame moving median filter. (C and D) Frequency of $Ca^{2+}$ signals in astrocyte somata (C) and neuropil (D) during overall wakefulness and overall sleep (*left*), during locomotion, whisking and quiet wakefulness (*middle*), and during NREM, IS and REM sleep compared to quiet wakefulness (*right*). (E and F) Duration of $Ca^{2+}$ signals in astrocyte somata (E) and neuropil (F) during overall wakefulness and overall sleep (*left*), during locomotion, whisking and quiet wakefulness (*middle*), and during NREM, IS and REM sleep compared to quiet wakefulness (*right*). (G and H) Same as in (E and F), but for amplitude of $Ca^{2+}$ signals instead of duration. All values represent estimates ± SEM, n = 6 mice, 284 trials for (C, E, G); 278 trials for (D, F, H). $^*p<0.05$, $^{**}p<0.005$, $^{***}p<0.0005$. For details on statistical analyses, see Methods.

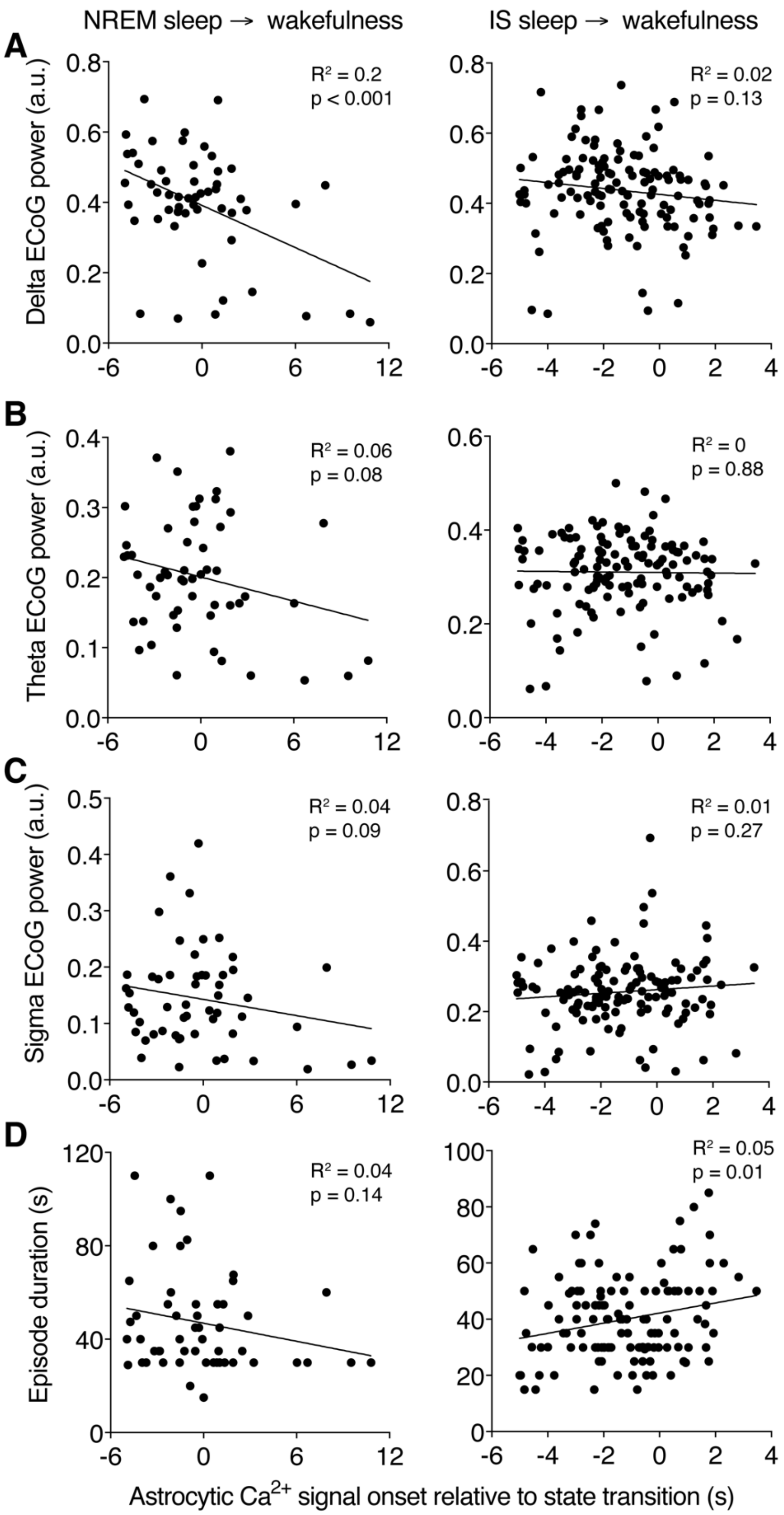


**Figure S9.** (A-D) Correlation between astrocytic $Ca^{2+}$ signal onset and ECoG power in delta (A), theta (B) and sigma (C) frequency bands, and NREM or IS sleep episode mean duration (D) in relation to state transition from NREM sleep (left) or IS sleep (right) to wakefulness. n = 6 mice, 59 NREM sleep to wakefulness transitions, 140 IS sleep to wakefulness transitions.

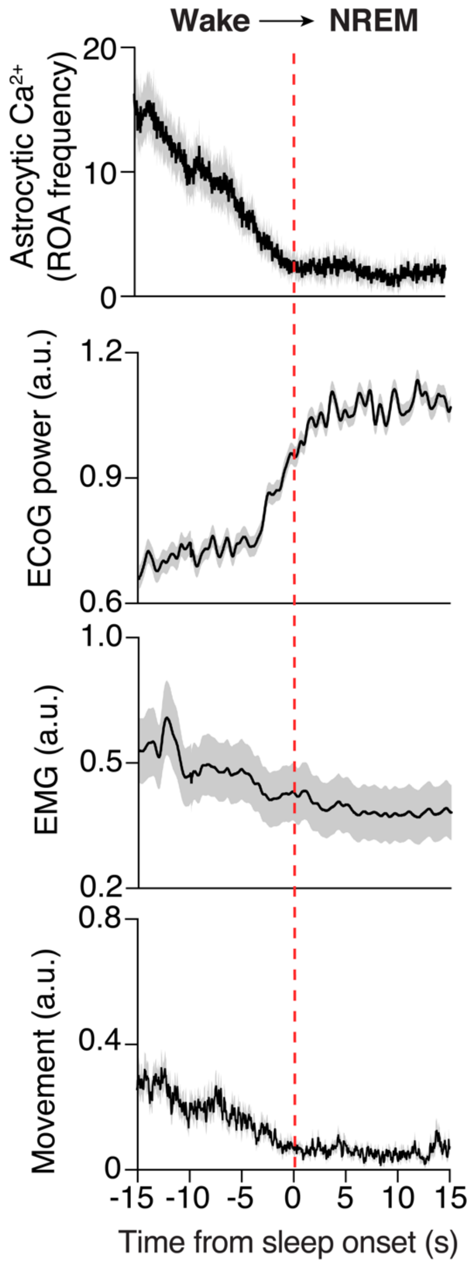


**Figure S10. Transition from wakefulness to sleep is associated with a decrease in astrocytic $Ca^{2+}$ signaling**. Mean time-course of (top to bottom) astrocytic $Ca^{2+}$ signals expressed as a number of ROAs per 100 $\mu m^2$ per minute, and traces of total ECoG power 0.5–30 Hz, EMG and mouse movement during transitions from wakefulness to NREM sleep. Data represented as mean ± SEM, n = 6 mice, 429 transitions.

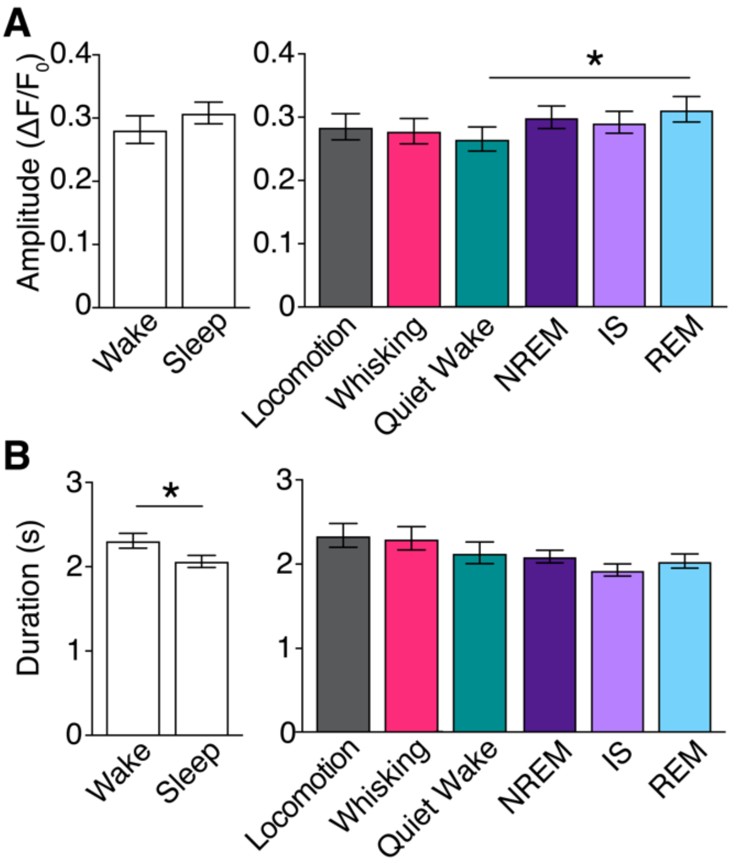


**Figure S11. $Ca^{2+}$ signaling in neuron somata across sleep-wake states.** (A–B) Amplitude (A) and duration (B) of $Ca^{2+}$ events in neuron somata ROIs in overall wakefulness and sleep (left) and across sleep-wake states (right). All values represent estimates ± SEM, n = 6 mice, 92 trials, *$p<0.05$, **$p<0.005$, ***$p<0.0005$. See also Figure 5. For details on statistical analyses, see Methods.

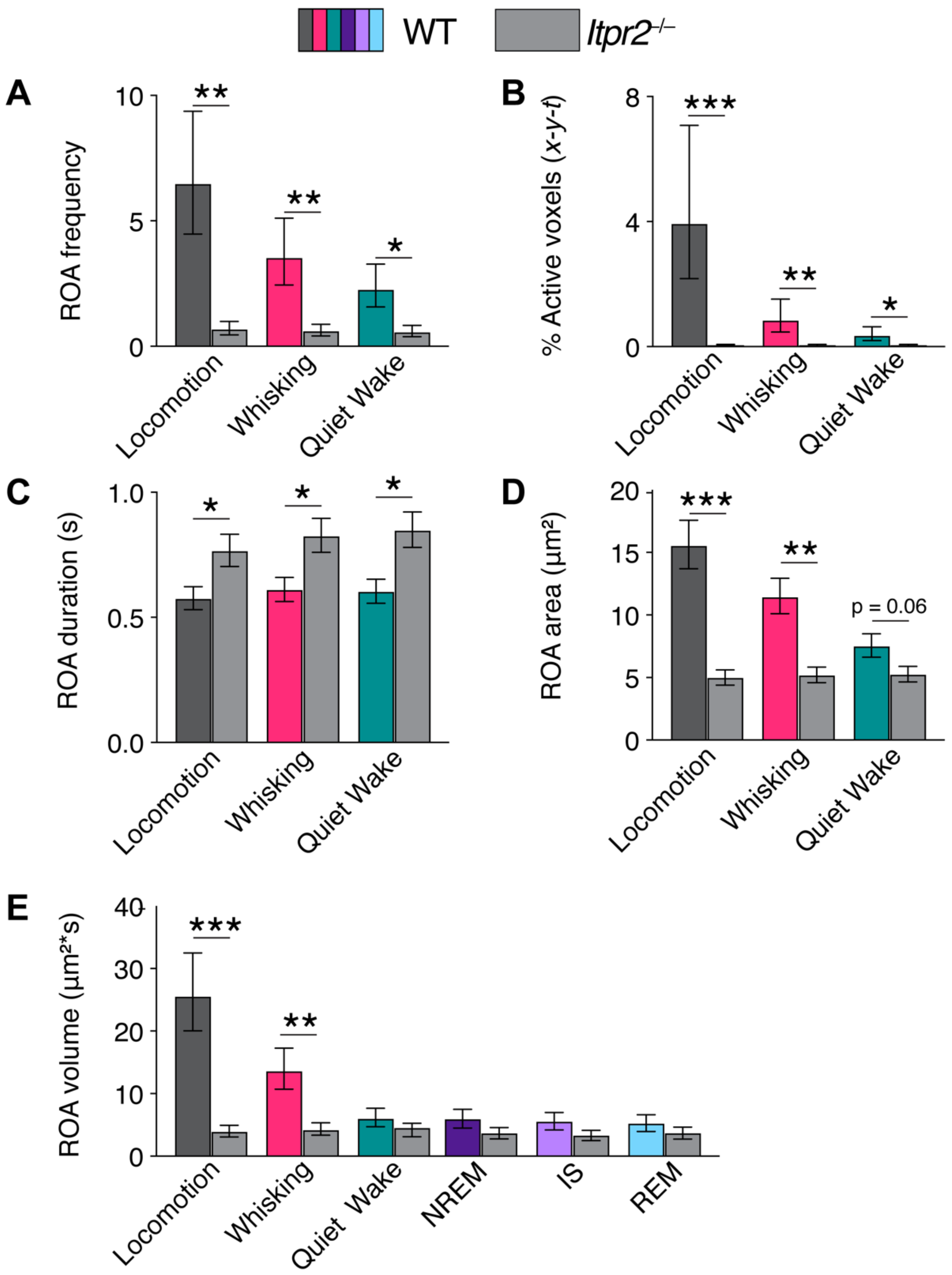


**Figure S12. Astrocytic $Ca^{2+}$ signaling in *Itrp2*$^{-/-}$ and WT mice.** (A to E) Mean ROA frequency expressed as number of ROAs per 100 µm$^2$ per minute (A), the percentage of active voxels (*x*-*y*-*t*) (B), ROA duration (s) (C), ROA area (µm$^2$) (D) during locomotion, whisking and quiet wakefulness, and ROA volume (µm$^2$*s) (E) across sleep-wake states in WT and *Itpr2*$^{-/-}$ mice. For WT mice: n = 6 mice, 82 trials; for *Itpr2*$^{-/-}$ mice: n = 6 mice, 96 trials. All values represent estimates ± SEM, *p<0.05, **p<0.005, ***p<0.0005. For details on statistical analyses, see Methods.

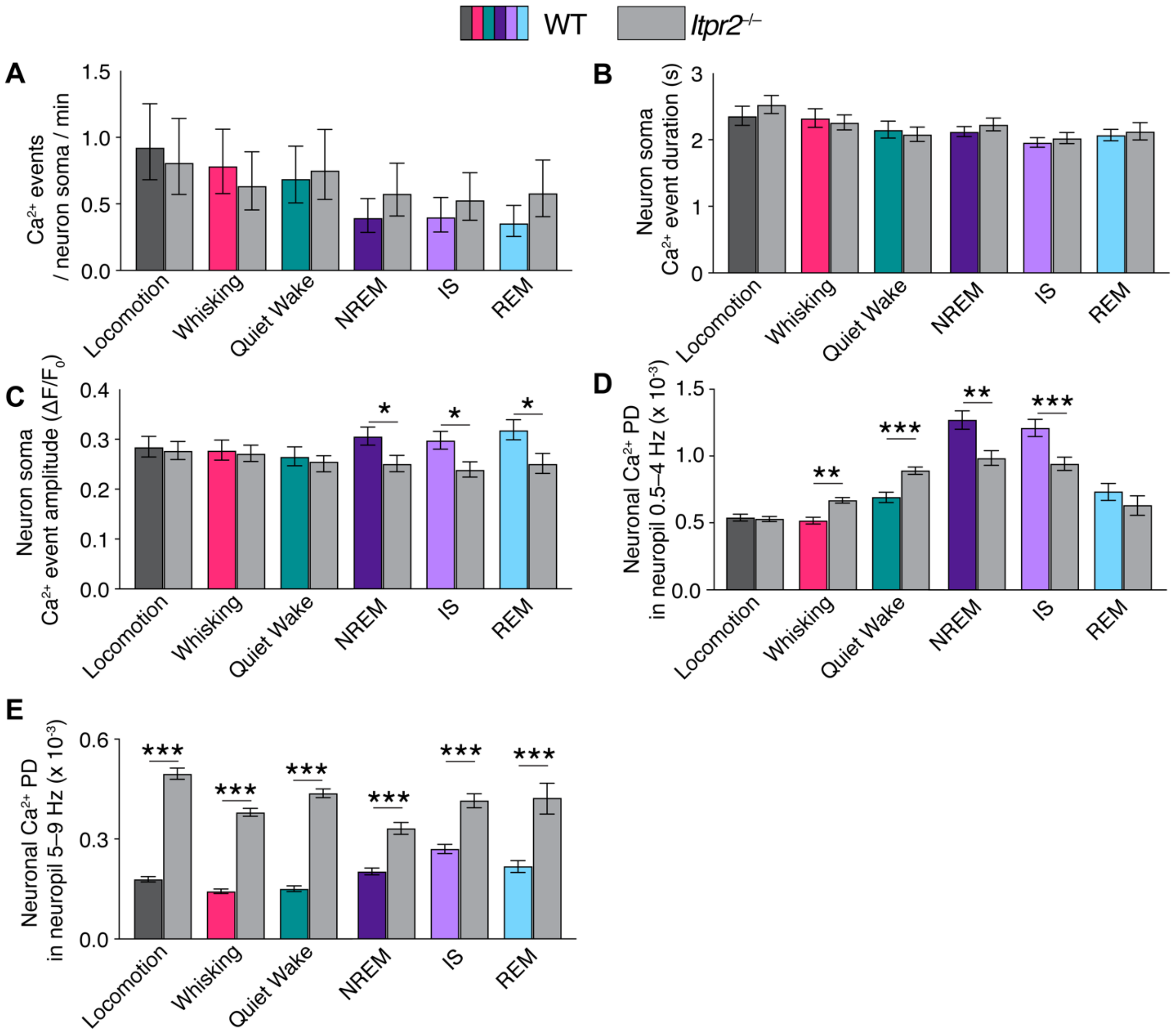


**Figure S13. Neuronal Ca$^{2+}$ signaling in *Itrp*2$^{-/-}$ and WT mice across sleep-wake states.** (A to C) Mean frequency expressed as events per minute per ROI (A), duration (B) and amplitude (C) of Ca$^{2+}$ signals in neuronal somata across sleep-wake states in WT and *Itpr2*$^{-/-}$ mice. For WT mice: n = 6 mice, n = 26 trials; for *Itpr2*$^{-/-}$ mice: n = 5 mice, n = 55 trials. (D and E) Power density (PD) of the neuronal Ca$^{2+}$ signal in neuropil in delta 0.5–4 Hz (D) and theta 5–9 Hz (E) frequency bands during across sleep-wake states in WT and *Itpr2*$^{-/-}$ mice. For WT mice: n = 6 mice, 25 trials; for *Itpr2*$^{-/-}$ mice: n = 5 mice, 70 trials. All values represent estimates ± SEM, *p<0.05, **p<0.005, ***p<0.0005. For details on statistical analyses, see Methods.
